# Design of a novel epitope-based tetravalent subunit vaccine against dengue virus: an immunoinformatics approach

**DOI:** 10.1101/2024.05.28.596168

**Authors:** Tanbin Jahan Ferdousy, Md Mehedy Hasan Miraz, Tawsif Al Arian, Md Afif Ullah, Md Tahsinul Haque Risat, Most. Afrin Akter, Shouhardyo Kundu, Bidduth Kumar Sarkar, Arghya Prosun Sarkar, Abul Bashar Ripon Khalipha, Sukalyan Kumar Kundu

## Abstract

Dengue imposes a profound global impact, with millions affected annually. Its transmission by *Aedes* mosquitoes poses significant challenges to combat, aggravated by urbanization and climate change. Despite efforts, no impeccable antiviral treatment exists to date, highlighting the urgency for a vaccine. Developing one encounters hurdles like the four distinctive serotypes of the virus and complex immune responses. In this study, we employed an immunoinformatics approach to design an epitope-based tetravalent subunit vaccine aimed at confronting all DENV serotypes. The study contemplates epitopes prediction, toxicity assessment, molecular docking, molecular dynamics (MD) simulations, immune simulations. On sequence retrieval, the epitopes were predicted and prioritized. The sequence of the finally designed vaccine was reached after a broad analysis of the antigenicity scores, serotype coverage, and population coverage. Human β-defensin 3 has been added as an adjuvant to the core vaccine sequence that comprises 23 epitopes and linkers. Notably, the vaccine has the highest antigenicity (0.9319) and 97.35% population coverage worldwide. The molecular docking operations of the vaccine with toll-like receptor 2 (TLR2) and TLR4 showed promising interactions with lowest energies of −1240.5 kJ/mol (76 members) and −1393.3 kJ/mol (40 members), respectively. Molecular dynamics (MD) simulations were run up to 200 nanoseconds, and the complexes of the vaccine with TLR2 and TLR4 were found to be very stable and flexible. Moreover, in immune simulations, the vaccine evoked robust immune responses. These findings suggest that our vaccine outperforms any other Dengue vaccine developed to date. However, since this study was conducted through *in silico* methods, *in vitro* and *in vivo* validations are required to confirm the vaccine as a potential candidate for clinical trials.

**Author summary:** We developed a novel epitope-based tetravalent subunit vaccine against all dengue virus (DENV) serotypes using immunoinformatics. Our vaccine demonstrated high antigenicity (0.9319) and wide population coverage (97.35%). Molecular docking and dynamics simulations indicated strong interactions with immune receptors which is crucial for the vaccines activity inside human body. Moreover, immune simulations showed robust responses indicating proper immunity against DENV. Therefore, our vaccine offers a promising solution to dengue fever, pending further *in vitro* and *in vivo* validations for clinical trials. Notably, this is the first ever dengue vaccine with such high efficacy in terms of immunoinformatics and vaccinomic approaches.

## Introduction

Dengue has posed a significant global health concern in recent decades. Around 4 billion individuals (nearly half of the global population) reside in regions vulnerable to dengue exposure [1]. Every year, roughly 400 million people incur dengue, and around 100 million get affected, whereas about 40,000 people die from severe manifestations [1]. The World Health Organization (WHO) documented more than 5 million dengue cases occurred with over 5,000 deaths in more than 80 territories and five WHO regions in 2023 [2]. Dengue is exposed in more than 100 countries in the American, Southeast Asian, Western Pacific, and African regions. The geographical escalation of dengue is evoked by climate change, urbanization, and increased global travel [3]. WHO has supported the use of integrated vector management, along with early and precise diagnosis, to lessen the impact of the disease. Despite these efforts, dengue continues to pose a challenge to public health systems worldwide and no impeccable cure has been established yet.

Dengue virus (DENV) is an RNA virus and a member of the *Flaviviridae* family. The virus is transmitted by mosquitoes of several *Aedes species*, mainly by *Aedes aegypti* [4]. There are four serotypes of the virus—DENV-1, DENV-2, DENV-3, and DENV-4—which are capable of producing independent diseases and even may exhibit immunological cross-reactivity with one another. An individual may experience all four serotypes of DENV in the life span. The viral genome encodes structural proteins (capsid protein, C; precursor membrane protein, prM; and envelope protein, E), and non-structural proteins (NS1, NS2a, NS2b, NS3, NS4a, NS4b, and NS5). The capsid (C) protein plays a crucial role in the packaging of the viral genome. It also interacts with the viral RNA during assembly and release [5]. The precursor membrane (prM) protein functions as a chaperone for the envelope (E) protein, thus, ensuring proper folding of the envelope protein and hereby enhancing the immunogenicity [6]. The protein E is essential for attachment and entry of the virus into host cells. It also induces neutralizing and protective antibodies and thus elicits the immune response of hosts [7].

Among the non-structural proteins, NS1 has various functions such as viral replication, evading the immune response by interfering with the host complement system, and affecting endothelial permeability to exacerbate disease severity [8]. NS2a has an important function in viral RNA replication, assembly of virions, and adjustment of the antiviral reaction of the host [9]. NS2b acts as a co-factor for the NS3 protease, playing a role in viral replication and potentially changing membrane permeability [10]. NS3 acts as a flexible protein necessary for copying the viral genome, performing tasks like RNA helicase and RNA triphosphatase activities, and being involved in viral polyprotein processing with its serine protease function [11]. It also impacts the immune responses of the host and aids in the development of viral pathogenesis. NS4a plays a role in creating the viral replication complex and altering membranes in host cells, which are important for RNA replication and avoiding the host’s antiviral defenses during DENV infection [12]. NS4b contributes to viral replication by engaging with additional nonstructural proteins, changing host cell responses to promote viral transmission, and inducing autophagy to form viral replication complexes and evade host immune responses [10]. NS5 acts as the RNA-dependent RNA polymerase necessary for replicating the viral genome, with its N-terminal region involved in processes like viral RNA capping and evading the immune system by blocking interferon signaling and altering host cell transcriptional responses in DENV infection [13].

The development of vaccines against DENV was initiated back in the 1920s but the dissimilitude among the four serotypes of DENV hindered the progress [14]. As of March 2024, only two dengue vaccines—Dengvaxia® and Qdenga®—are commercially available and several vaccines are at different phases of clinical trials [15, 16]. Despite being tetravalent, a major issue with Dengvaxia® is its efficacy is limited to only individuals ages 9-16 years who had been affected with DENV before getting vaccinated with this vaccine [17]. Dengvaxia® has also been reported to increase the risk of severe dengue in people having no prior dengue infections [17, 18]. In contrast, Qdenga® is not limited to such conditions being applicable to anyone previously infected or not [19]. Another study shows that TAK-003—later known as Qdenga®—in phase 3 clinical trial, exhibited efficacy for long term and protection against all four DENV serotypes in individuals exposed to the virus previously [20]. DENV-1 and DENV-2 were successfully safeguarded by TAK-003 in dengue-naive individuals. This vaccine, however, has not been approved yet by the Food and Drug Administration (FDA).

This study approached a reverse vaccinomic strategy based on immunoinformatics to design a tetravalent subunit dengue vaccine with multiple epitopes derived from all serotypes of DENV. A key advantage in designing multi-epitope vaccines is the absence of live components thereby deteriorating disease risks. The vaccines can be produced at the laboratory level without the involvement of any pathogenic components or organisms. By incorporating technologies for protein production and purification, the protein can be conveniently synthesized and purified. Due to less tendency of allergic reactions, cost-efficacy, and rapid development, these approaches are preferred over traditional vaccine development methods. Similar approaches have been taken previously in developing vaccines against Zika, Ebola, chikungunya, SARS-CoV-2, and even DENV, but no vaccine could come into the commercial market [21–25]. Notably, our designed vaccine surpassed all dengue vaccines in terms of antigenicity, population coverage, serotype coverage, structural integrity, and many other aspects.

While designing the vaccine, B-cell and T-cell epitopes were taken into consideration which were adjuvanted with human β-defensin 3 at the N-terminal to elicit proper immune responses. Upon assembling the sequence, structural validations, physicochemical properties, and solubility were assessed for the initial scrutiny of the vaccine. Molecular docking approaches were conducted to confirm the extent of the interaction of the vaccine with the immune receptors of the body. Additionally, molecular dynamics (MD) simulations were run to study the stability, flexibility, and further parameters of the interactions. Immune simulations were run to observe the immune response of the human body in response to vaccination. Finally, the vaccine was cloned inside an expression vector using an *in silico* cloning approach.

## Methodology

### Viral protein sequence retrieval

The initial step for designing the vaccine was the retrieval of the viral protein sequence of DENV-1 to −4 (Fig 1). The complete viral protein sequences of all dengue serotypes were selected for designing the tetravalent subunit vaccine. The polyprotein sequences from the complete genome of DENV-1 (Accession: NC_001477), DENV-2 (Accession: NC_001474), DENV-3 (Accession: NC_001475), and DENV-4 (Accession: NC_002640) were retrieved in FASTA format from the NCBI Genbank (https://www.ncbi.nlm.nih.gov/genbank/) [26]. Afterwards, the sequences were analyzed on basis of all structural and functional proteins of dengue virus. The antigenicity of each protein of each sequence were determined from VaxiJen 2.0 server (https://www.ddg-pharmfac.net/vaxijen/VaxiJen/VaxiJen.html) [27]. While submitting to VaxiJen, the target organism was set to virus and the threshold value was set to 0.4. The antigenicity score was noted and compared for further analysis.

**Fig 1.**
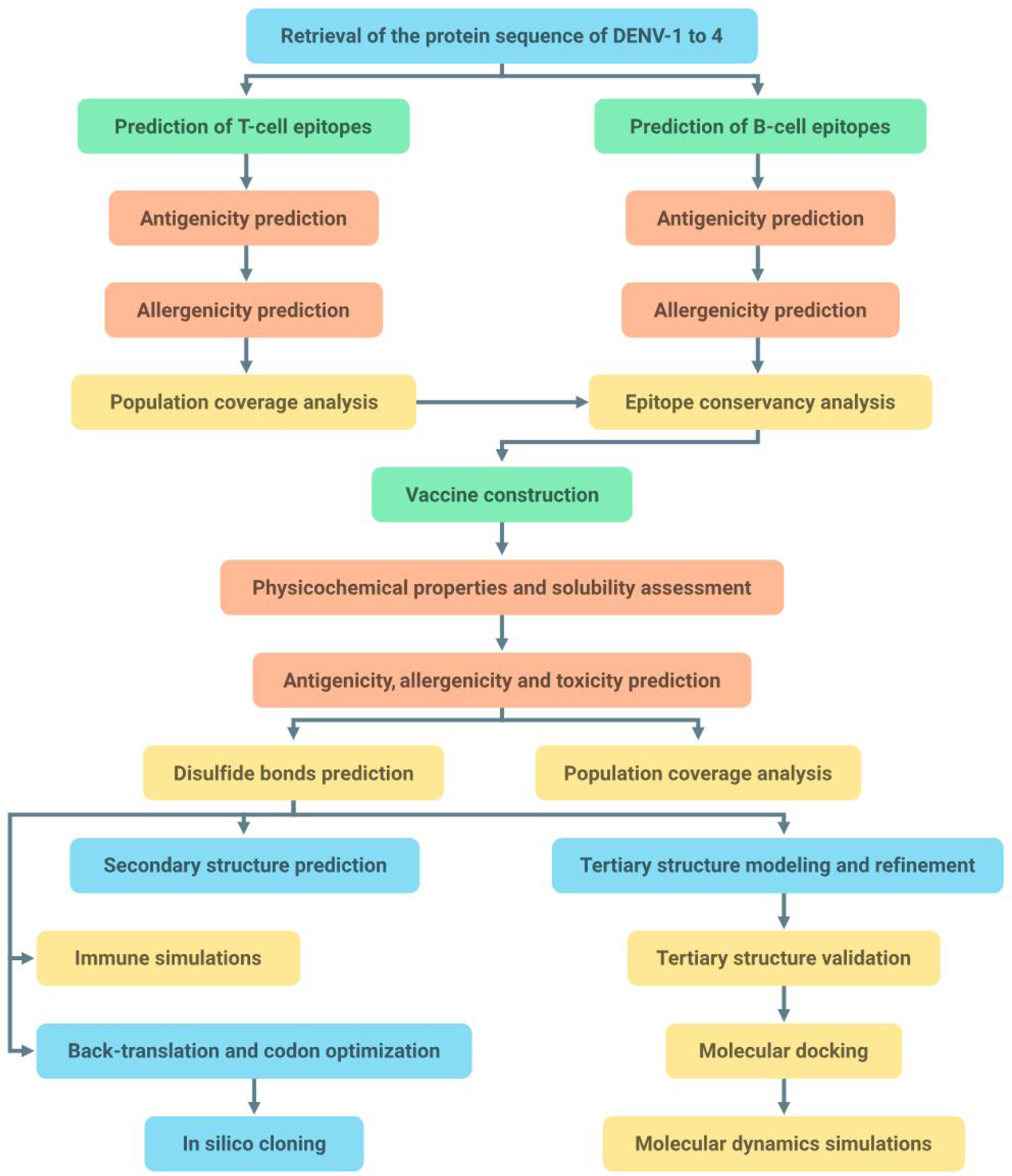
The workflow of the DENV vaccine design.

### B-cell epitopes prediction

B-cell epitopes were predicted from ABCpred (https://webs.iiitd.edu.in/raghava/abcpred/) [28]. The retrieved sequences for all serotypes in one letter-code without any header were pasted on this server individually. The prediction threshold value was set to 0.83, and the length of each epitope was fixed to 16 amino acids for each submission. Before submission of the sequences for prediction, the overlapping filter was activated. The ABCpred server employs an artificial neural network trained on linear B-cell epitopes obtained from the Bcipep database (https://webs.iiitd.edu.in/raghava/bcipep) for epitope prediction [28]. The maximum accuracy achieved was through a recurrent neural network (Jordan network) comprising a single hidden layer with 35 hidden units, using a window length of 16. Through fivefold cross-validation, the final network demonstrated an overall prediction accuracy of 65.93%, whereas the sensitivity, specificity, and positive prediction values were found to be 67.14%, 64.71%, and 65.61%, respectively [28].

### T-cell epitopes prediction

The cytotoxic T-lymphocyte (CTL) epitopes and helper T-lymphocyte (HTL) epitopes were predicted as MHC class I and class II binding epitopes, respectively, from the IEDB Immune Epitopes Database & Tools server (https://www.iedb.org/) [29]. During MHC-I binding prediction, the FASTA sequences were pasted individually and the prediction method was set to ANN 4.0. The whole HLA allele reference set was selected from the list. Afterwards, the epitopes were sorted based on IC_50_ values and epitopes having IC_50_ < 100 were prioritized for further analysis. In contrast, MHC-II binding epitopes were predicted using NN-align 2.3 (NetMHCII 2.3) prediction method for the full HLA reference set. The MHC-I binding epitopes were predicted in 9-meric form, whereas MHC-II epitopes were set to 15-mers.

### Antigenicity and allergenicity prediction of the epitopes

Antigenic property of the derived B-cell and T-cell epitopes were predicted using the VaxiJen v2.0 server [27]. The prediction of antigens in VaxiJen is alignment-free and relies on a range of physicochemical properties inherent to proteins [27]. While submitting each epitopes for prediction, the target organism was selected as virus and the threshold value was set to 0.4.

All the epitopes surpassing the threshold value were considered as probable antigen and thus were subjected for allergenicity prediction. To predict whether these epitopes were non-allergen, AllerTOP v.2 server (https://www.ddg-pharmfac.net/AllerTOP/) was used [30]. The AllerTOP v.2 web server uses the auto cross covariance (ACC) method to convert protein sequences into uniform vectors. ACC, developed by Wold et al. (1993), employs five E descriptors from Venkatarajan and Braun (2001) to represent amino acid properties [31, 32]. These descriptors include hydrophobicity, molecular size, helix-forming propensity, relative abundance, and β-strand forming propensity. Classification is done using k-nearest neighbor (kNN, k=1) on a training set of 2427 known allergens and an equal number of non-allergens. Once the prediction for all epitopes were done, allergens were removed, and non-allergens were shortlisted.

### Conservancy and population coverage analysis of epitopes

The conservancy of each epitopes were analyzed using the Epitope Conservancy Analysis tool of IEDB Analysis Resource (http://tools.iedb.org/conservancy/) [29, 33]. This analysis was performed to scrutinize how much the epitopes were conserved in the 4 serotypes of dengue. The threshold was set to a minimum of 80% while submission for analysis. Since the vaccine needs to be tetravalent to combat all serotypes, the epitopes conserved in the maximum number of serotypes were preferred. Afterwards, Population Coverage server of IEDB Analysis Resource (http://tools.iedb.org/population/) was utilized to detect the population coverage of each MHC-I and -II epitope classes [29]. The MHC-I and -II epitopes were uploaded to this server along with their respective alleles. The prediction was queried by “area_country_ethnicity” among the world population. The detailed tabulations from the server were analyzed for each MHC classes and epitopes were prioritized based on the worldwide coverage.

### Vaccine construction

An epitope-based subunit vaccine requires three types of elements—adjuvant, epitopes, and linkers—to elicit proper immune response as a complete vaccine. The vaccine sequence commences with the adjuvant when integrated into the primary peptide chain of the vaccine. The adjuvant plays a crucial role in stimulating the desired antigen-specific immune response, thereby enhancing the effectiveness of vaccination [34]. Human β-defensin 3 was included in this vaccine as the adjuvant due to its dual functionality as both an antimicrobial agent and an immunomodulator [35]. The FASTA sequence of human β-defensin 3 (PDB ID: 1KJ6) was retrieved from the RCSB Protein Data Bank [36–38]. The adjuvant was added to the main vaccine sequence with a EAAAK linker. Subsequently, the MHC-I epitopes were included adjacent to the adjuvant and were connected to each other using AAY linkers. Next, the MHC-II epitopes were added, linked together by GPGPG linkers. Finally, the B-cell epitopes were incorporated into the sequence, with each epitope connected by KK linkers. This sequential arrangement ensures proper spacing and connectivity between the adjuvant, epitopes, and linkers, facilitating the induction of an effective immune response.

### Physicochemical properties and solubility assessment

The PepCheck (https://lab.oimi.co/pepcheck/) and ExPASy ProtParam (https://web.expasy.org/protparam/) tools were employed to evaluate the physicochemical properties of the vaccine [39]. The amino acid sequence was entered into the server for assessment. Various parameters including the number of amino acids, molecular weight, molecular formula, theoretical isoelectric point (pI), amino acid compositions, extinction coefficients, estimated half-life, instability index, aliphatic index, and grand average of hydropathicity (GRAVY) values were analyzed in detail. Afterwards, the solubility of the vaccine construct was assessed and validated using three servers—SOLpro from SCRATCH Protein Predictor (https://scratch.proteomics.ics.uci.edu/) [40, 41], SoluProt v1.0 [42], and Protein-Sol [43]. SOLpro utilizes a two-stage SVM architecture to predict the propensity of proteins to be soluble upon overexpression in *E. coli*. This method is based on multiple representations of the primary sequence [41]. Each classifier in the first layer takes a distinct set of features describing the sequence as input. Subsequently, a final SVM classifier aggregates the predictions from the first layer classifiers to predict whether the protein is soluble, along with providing a probability estimate. Similarly, SoluProt v1.0 also predicts the solubility of proteins upon expression in *E. coli* [42].

### Antigenicity, allergenicity and toxicity prediction of the vaccine

The antigenicity of the designed vaccine was predicted from the VaxiJen v2.0 and was validated using ANTIGENpro from SCRATCH Protein Predictor [41]. The FASTA sequence of the vaccine construct was uploaded to VaxiJen. The target organism was chosen as virus keeping the threshold value a minimum of 0.4. After retrieval of the antigenicity from VaxiJen, it was further validated from the ANTIGENpro tool. Consequently, the vaccine was subjected to allergenicity and toxicity prediction. The AllerTOP v.2 server was utilized to predict whether the vaccine construct is non-allergenic. Afterwards, the FASTA format of the sequence was submitted to the T3DB server (http://www.t3db.ca/) for toxicity prediction [44]. In this process, the costs to open and extend a gap were set to −1, the penalty for mismatch was set to −3, and the reward for match was set to 1. The expectation value was set to 0.00001. Gapped alignment was performed, and the query sequence was filtered by checking the respective checkboxes during sequence submission. Finally, by conducting the search, the toxicity of the vaccine was predicted.

### Population coverage analysis of the vaccine

Population Coverage server was again used at this step to assess the population coverage of the vaccine construct [29]. The two epitope classes (MHC-I and MHC-II) were uploaded with their respective alleles as done before. The prediction was queried also as before by selecting “area_country_ethnicity” among the all population of the world. The calculation was conducted for each MHC classes separately, and then combined together. The coverage, average hit, and PC90 values were scrutinized. The standard deviation was also taken into account while analyzing.

### Disulfide bonds prediction

The prediction of disulfide bonds in the vaccine construct was performed using the DIpro tool of the SCRATCH Protein Predictor server. DIpro is a cysteine disulfide bond predictor that utilizes a 2D recurrent neural network. It supports various algorithms including SVM, graph matching, and regression [45]. This tool is capable of predicting whether a peptide sequence contains disulfide bonds, estimating the number of disulfide bonds present, and predicting the bonding state of each cysteine and the bonded pairs. Upon submission, the peptide sequence undergoes processing in two steps [45]. Firstly, DIpro employs SVM to classify whether the sequence contains disulfide bonds. Subsequently, it utilizes neural network and graph algorithms to predict the number of disulfide bonds and the bond pattern.

### Secondary structure prediction

Secondary structure of the vaccine was predicted from the PSIPRED 4.0 server (http://bioinf.cs.ucl.ac.uk/psipred/) [46]. Upon submission of the protein sequence, PSIPRED generated three types of sequence plots and a cartoon. The figures were analyzed to predict the structure of different portions in the vaccine peptide chain as strand, helix, or coil. The type of amino acids—small nonpolar, hydrophobic, polar, or aromatics plus cystiene—were also obtained from a plot. Moreover, from the cartoon, the structures were obtained with different confidence levels of prediction.

### Tertiary structure modeling and refinement

The tertiary structure of the vaccine was predicted using the 3Dpro tool of the SCRATCH Protein Predictor server. The peptide sequence of the vaccine was pasted to the server and submitted for the prediction. 3Dpro utilizes predicted structural features and PDB knowledge-based statistical terms to construct multiple peptide models using random seeds. The model with the lowest energy is selected as the final prediction. Notably, this method is currently a de novo approach, meaning it does not rely on structural templates. Hence, the PDB file of the vaccine 3D model was retrieved from 3Dpro results. Following the initial modeling, the structure was refined using the GalaxyRefine tool from the GalaxyWEB server (https://galaxy.seoklab.org/cgi-bin/submit.cgi?type=REFINE) [47–49]. The PDB file of the tertiary structure was uploaded and submitted for refinement. From the refinement results, the GDT-HA, MolProbity, and Rama favored scores were obtained. The model with the highest Rama favored score was chosen for further validation.

### Tertiary structure validation

The Ramachandran plot of the peptide sequence was analyzed using the PROCHECK tool of the SAVES v6.0 structure validation server (https://saves.mbi.ucla.edu/) [50]. The tertiary refined model was uploaded to the server, and the PROCHECK tool was executed. The Ramachandran plot provided information on the allowed regions of each amino acid residue, aiding in the interpretation of the structural quality. Additionally, chi1-chi2 plots, main-chain and side-chain parameters, residue properties, G-factors, and planar groups were analyzed to further assess the structural properties of the model. Afterwards, the Z-score of the peptide model was evaluated using ProSA-web (https://prosa.services.came.sbg.ac.at/prosa.php) [51]. The Z-score provides an indication of the overall quality and compatibility of the model with experimental data and serves as a measure of its structural integrity (Wiederstein and Sippl, 2007).

### Molecular docking of the vaccine with immune receptors

Molecular docking was carried out to investigate the interaction between the vaccine construct and immune receptors in the human body. The docking tasks were performed using the ClusPro 2.0 server (https://cluspro.bu.edu/) on supercomputers [52–55]. The vaccine model was docked separately with TLR2 (PDB ID: 2Z7X) and TLR4 (PDB ID: 3FXI), whose PDB structures were obtained from the RCSB PDB server [56, 57]. To ensure efficient processing, both the CPU and GPU servers of ClusPro were utilized to run and validate the docking results. ClusPro employs FFT-based global sampling using PIPER to explore docking orientations efficiently, followed by clustering to identify low-energy conformations [54]. It then utilizes CHARMM minimization to refine the structures, removing steric clashes. The interactions of the docked complexes were scrutinized using the PDBsum tool (https://www.ebi.ac.uk/thornton-srv/databases/pdbsum/) [58].

### Molecular dynamics simulations

Molecular dynamics is a sophisticated automated simulation approach used for evaluating the degree of stability of protein and protein-ligand complex structure at the minuscule stage through demonstrating the behavioral characteristics, interacting arrangement, fluctuation, the physical foundation of function, and structure. The molecular dynamics (MD) simulations were conducted using GROMACS version 2023.1 to investigate the behavior of both vaccine-TLR4 complex and vaccine-TLR2 complex [59]. The protein was parameterized for its protein content using the CHARMM General Force Field. The SwissParam server was utilized to perform the ligand topologies [60]. The structures underwent 2500 cycles of vacuum minimization using the steepest descent method in order to mitigate any potential steric issues. The solvation of the structure was accomplished through the utilization of the Simple Point Charge (SPC) water model. Subsequently, the system was rendered neutral by introducing Na^+^ and Cl^-^ ions utilizing the gmx genion tool. This measure was implemented in order to maintain the overall electrical neutrality of the system. After minimization, three steps were conducted in the MD simulation: NVT, NPT, and production. The equilibration of the systems was conducted in two phases. Initially, an NVT equilibration lasting 100 picoseconds was conducted to attain a steady state of the number of particles, volume, and temperature. The purpose of this step was to elevate the system to a temperature of 300 Kelvin. The second step involved conducting a 100 picoseconds NPT equilibration, which aimed to maintain the system’s pressure and density stability by ensuring an equal number of particles, pressure, and temperature. The simulations involved the imposition of bond constraints on all bonds within the protein, thereby inducing position restraint of the protein group. The constrained conditions of NVT and NPT resulted in the relaxation of water molecules surrounding the protein, leading to a decrease in system entropy. The dynamics were conducted utilizing the Parrinello-Rahman barostat and the v-rescale thermostat [61]. The relaxation of the barostat and thermostat persisted for a duration of 100 picoseconds. The application of Linear Constraint Solver algorithm was utilized to impose constraints on the covalent bonds. The Particle-Mesh Ewald (PME) method was employed to process the electrostatic interactions. After reaching equilibrium, every system underwent a production run that lasted for 50 nanoseconds (ns) of simulation time.

### Immune simulations

Immune simulations were run to scrutinize the immune response inside the body. C-ImmSim server (https://kraken.iac.rm.cnr.it/C-IMMSIM/) was used to run the immune simulations [62–64]. The simulations were run under a random seed. The simulation volume was set to 10 and the simulation was run for 1000 steps (for around 333 days). The host HLA selections were chosen for A0101, A0201, B0702, B0801, DRB1_0101, and DRB3_0101 respectively. Total 3 injections were included at the interval of 20 days. The time steps with this interval represents 1, 61, and 121, respectively, for the 1st, 21st, and 41st days. The number of antigens for each dose was set to 1000. The FASTA sequence of the vaccine was inputted as the Ag molecule. After setting up the pre-submission parameters, the vaccine was submitted for the simulations.

### Back-translation, optimization and *in silico* cloning

Back-translation of the vaccine peptide sequence was conducted using the EMBOSS Backtranseq server (https://www.ebi.ac.uk/Tools/st/emboss_backtranseq/), selecting the *Esh coli* K12 strain for codon usage [64]. The resulting back-translated sequence was retrieved from the server. Later on, Java Codon Adaptation Tool (JCAT; https://www.jcat.de/) was utilized for codon optimization, with the organism set as *E. coli* (strain K12) [66]. During optimization, avoidance of rho-independent transcription terminators, prokaryotic ribosome binding sites, and *Eco*RI restriction enzyme cleavage sites was ensured. The codon adaptation index (CAI) and the GC-content of the improved sequence were recorded. A higher CAI value (0.8–1.0) is always expected as higher CAI indicates higher expression [67]. Afterwards, *in silico* restriction cloning of the vaccine was performed using SnapGene software (www.snapgene.com), employing the *E. coli* pET-28a(+) vector. Prior to cloning, *Nde*I and *Xba*I restriction enzyme sites were incorporated at the N- and C-terminals of the optimized DNA sequence. Finally, the DNA sequence was inserted into the pET-28a(+) vector.

## Results

### Viral protein sequence

The viral polyprotein sequences of DENV-1, DENV-2, DENV-3, and DENV-4 were 3392, 3391, 3390, and 3387 amino acids in lengths, respectively (<u>S1 File</u>). The polyprotein comprises of structural and non-structural proteins. The structural proteins of the virus play a pivotal role in facilitating host invasion and orchestrating the intricate assembly of viral particles. In contrast, non-structural proteins assume a key function in the viral life cycle, secreting diverse enzymes that actively contribute to the intricate processes of viral replication and the synthesis of structural proteins. The interplay between these structural and non-structural components are crucial in the viral life cycle.

### B-cell epitopes prediction and prioritization

B-cell epitopes were predicted for the complete sequence of all DENV serotypes utilizing the ABCPred algorithm. The shortlisting of epitopes hinged upon a stringent criterion, incorporating a prediction score ≥ 0.83, alongside considerations of antigenicity, allergenicity, and toxicity. Only probable antigens and probable non-allergens were sorted out for further progress. The length of each B-cell epitopes were 16 amino acids as specified while submitting the sequence for the prediction. Moreover, the B-cell epitopes that were found in the maximum serotypes of DENV were favored. Among the selected epitopes, DENV-2 was represented by only one B-cell epitope, whereas all other serotypes had 2 epitopes for each, of which some are common. The finally selected B-cell epitopes are presented in Table 1.

**Table 1.** Prioritized B-cell epitopes, their antigenicity, and conservancy across different DENV serotypes.

| B-cell Epitopes | Antigenicity Score | DENV-1 | DENV-2 | DENV-3 | DENV-4 |
| --- | --- | --- | --- | --- | --- |
| RNLTIMDLHPGSGKTR | 1.5229 | ✓ |  | ✓ |  |
| TAGWDTRITEDDLQNE | 0.7913 | ✓ |  |  | ✓ |
| APSYGMRCVGVGNRDF | 1.9706 |  | ✓ |  |  |
| KTKKDLGLGSIATQQP | 1.4308 |  |  | ✓ |  |
| TGEIGAIALDFKPGTS | 1.7277 |  |  |  | ✓ |

### T-cell epitopes prediction and prioritization

T-cell epitopes, crucial for immune response, were predicted from the IEDB T-cell epitope prediction tools, discerning between MHC-I (9-mer) and MHC-II (15-mer) epitopes. Several factors were assessed while prioritizing the epitopes: combined score, reflecting overall epitope quality; antigenicity, indicating the likelihood of eliciting an immune response; allergenicity, to avoid potential allergic reactions; and IC_50_ values below 100 nM, denoting strong binding affinity to MHC molecules. The epitopes were then subjected to antigenicity and allergenicity prediction, and the epitopes with higher antigenicity scores and probable non-allergenic properties were prioritized. Moreover, epitope conservancy across DENV serotypes was scrutinized for maximum DENV serotypes coverage. For ensuring common antigenic regions across different DENV strains, epitopes shared among all or most serotypes were kept at top priority. The finally selected T-cell epitopes of both MHC-I and -II classes are shown in Table 2.

**Table 2.**
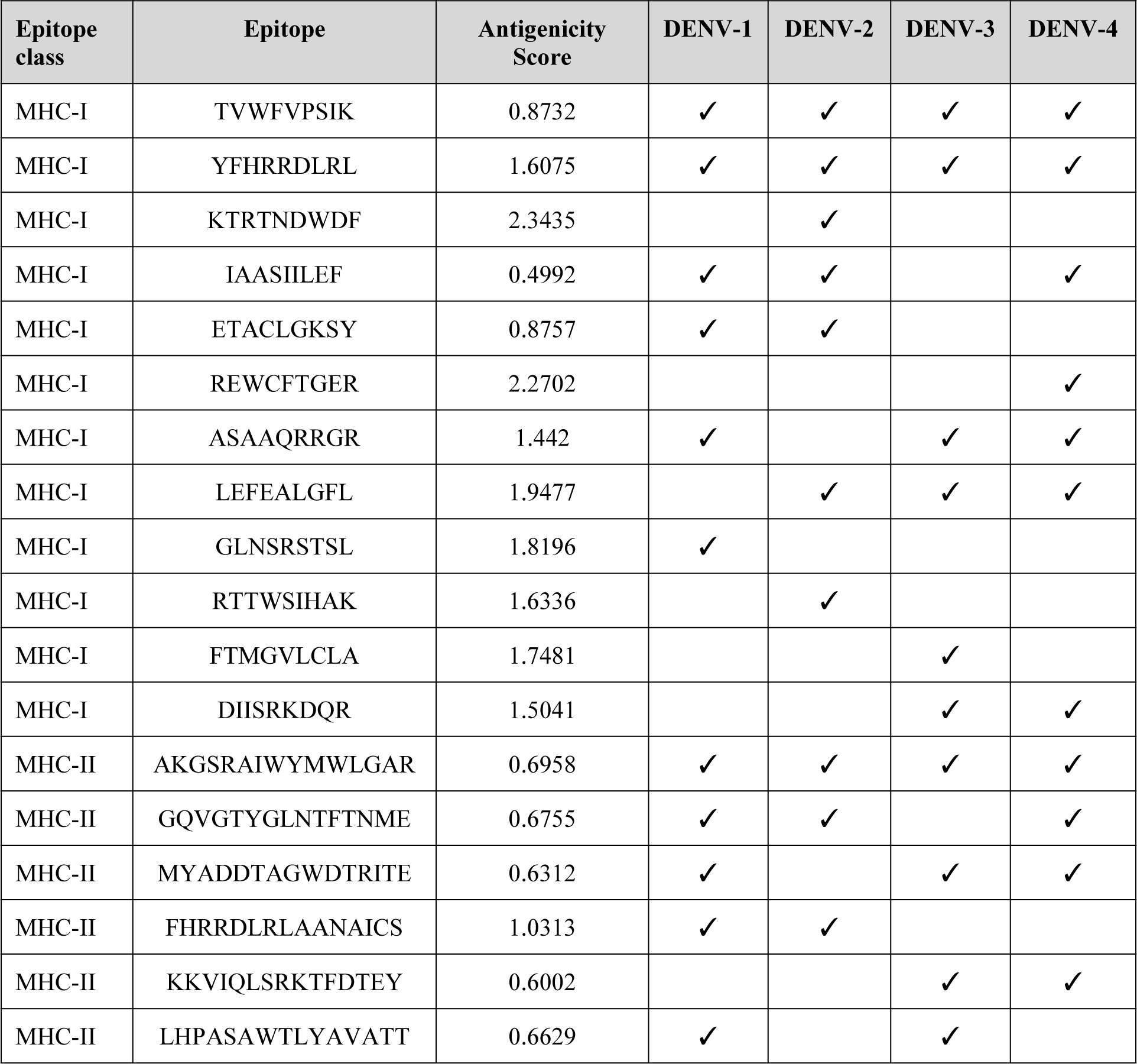
Prioritized T-cell epitopes, their antigenicity, and conservancy across different DENV serotypes. The epitopes are shown based on classes (MHC-I and -II).

### Vaccine construct

The final vaccine exhibited a length of 401 amino acids. The N-terminal residue of the construct was Glycine (Gly). The adjuvant human β-defensin 3 was linked to the core vaccine sequence with an EAAAK linker. The vaccine construct encompassed a total of 23 epitopes, comprising 12 MHC-I epitopes, 6 MHC-II epitopes, and 5 B-cell epitopes (Fig 2). Among these, DENV-1, DENV-2, DENV-3, and DENV-4 were represented by 13, 11, 12, and 13 epitopes, respectively. Moreover, among the 12 MHC-I epitopes, 2 were found common across all DENV serotypes. To be more specific, 6 MHC-I epitopes were attributed to DENV-1, 7 to DENV-2, 6 to DENV-3, and 7 to DENV-4. In contrast, among the MHC-II epitopes, DENV-1, DENV-2, DENV-3, and DENV-4 were specific to 5, 3, 4, and 4 epitopes, respectively, with a subset being common across the serotypes.

**Fig 2.**
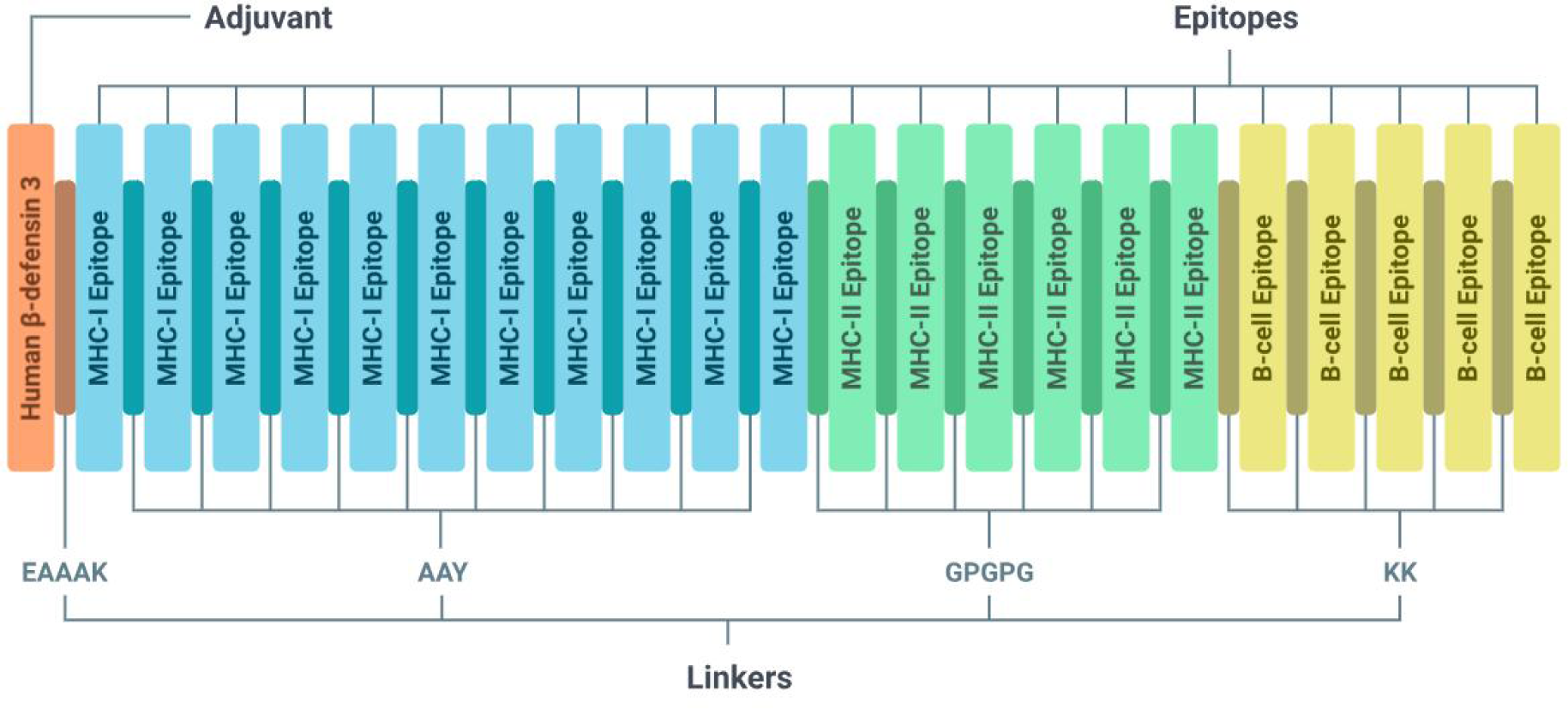
Vaccine construct. The adjuvant, 3 types of epitopes, and 4 types of linkers are shown in different colors.

### Physicochemical properties and solubility assessment

Physicochemical properties of the construct were assessed from PepCheck (https://lab.oimi.co/pepcheck/) and thus were validated using ExPASy ProtParam tool. The molecular formula of the vaccine was developed as C_1950_H_3053_N_525_O_555_S_17_ and the molecular weight stood at 43.84 kDa. The major parameters, involving molecular weight, atom count, total negatively charged residues (Asp + Glu), total positively charged residues (Arg + Lys), and isoelectric point (pI), which fell within optimum ranges (Table 3). Stability assessment, characterized by an instability index below 40, indicated the vaccine’s robust nature. The vaccine, with a GRAVY value below 0.00, exhibited hydrophilic properties. Solubility predictions from SOLpro, SoluProt 1.0, and Protein-Sol scored 0.823, 0.794, and 0.519, respectively. The scores affirmed the soluble nature of the vaccine construct.

**Table 3.** Physicochemical properties and solubility of the vaccine construct.

| Parameters | Vaccine construct measures |
| --- | --- |
| Molecular weight | 43.84 kDa |
| Number of atoms | 6140 |
| Number of amino acids | 401 |
| Total number of negatively charged residues (Asp + Glu) | 32 |
| Total number of positively charged residues (Arg + Lys) | 63 |
| Theoretical isoelectric point (pI) | 9.81 |
| Extinction coefficient | 81415 M<sup>-1</sup>cm<sup>-1</sup> |
| Instability index | 28.06 |
| Estimated half life | 30 hours (mammalian reticulocytes, *in vitro*) |
|  | >20 hours (yeast, *in vivo*) |
|  | >10 hours (*Escherichia coli*, *in vivo*) |
| Aliphatic index | 64.94 |
| Grand average of hydropathicity (GRAVY) | -0.444 |
| Solubility | 0.823 (SOLpro) |
|  | 0.794 (SoluProt 1.0) |
|  | 0.519 (Protein-Sol) |

### Antigenicity, allergenicity and toxicity of the vaccine

The antigenicity of the vaccine was 0.9319 as per VaxiJen 2.0 which is significantly higher than the threshold 0.4. The antigenicity was verified using ANTIGENpro server and it was 0.813649. Afterwards, on assessment of the allergenicity of the vaccine, the vaccine was found to be probable non-allergen as per AllerTOP v.2 and AlgPred server. AlgPred showed a score of −0.63394281 for the vaccine candidate, that surpasses the threshold of maximum −0.40 to be non-allergen. Finally, on submission of the FASTA sequence of the vaccine to the T3DB server, no results came from the database, thus indicating the vaccine as non-toxic.

### Population coverage

The worldwide population coverage of the vaccine was 97.35% as per IEDB Population Coverage server (<u>S1 Fig</u>). Such extensive coverage indicates the potential for widespread immunization and highlights the vaccine’s substantial impact in combating the targeted pathogen. The average hit combining both MHC-I and -II epitopes was 3.53 with a PC90 of 1.74. For MHC-I, 85.42% population coverage was obtained with average hits of 2.02 and PC90 of 0.69. In contrast, for MHC-II, the population coverage was 81.81% with average hits of 1.50 and PC90 of 0.55. The average hits are the average number of epitope hits or, HLA combinations recognized by the population, whereas the PC90 value is the minimum number of epitope hits or, HLA combinations recognized by 90% of the population.

### Disulfide bonds analysis

DIpro tool of SCRATCH Protein Predictor predicted that the sequence does have disulfide bonds. This tool detected 11 cysteines in the amino acid chain and predicted 4 disulfide bonds that may form in the sequence. Specifically, cysteines at positions 11, 18, 33, 40, 102, 114, 270, and 357 are anticipated to participate in the formation of these disulfide bonds. The classification results were obtained using Support Vector Machine (SVM). The predicted bonds (cysteine pairs) are listed in Table 4, ordered by probability in descending order.

**Table 4:** Predicted disulfide bonds (cysteine pairs). The bonds shown in the table are ordered by probability in descending order.

| Bond Index | Cys1 Position | Cys2 Position |
| --- | --- | --- |
| 1 | 11 | 18 |
| 2 | 270 | 357 |
| 3 | 102 | 114 |
| 4 | 33 | 40 |

### Secondary structure

The PSIPRED 4.0 server generated the secondary structure prediction for the vaccine peptide based on the submitted amino acid sequence. In the provided output, pink regions represent helices, yellow regions indicate strands, and gray regions resemble coils (Fig 3). Additionally, the confidence levels are denoted by the darkness of the color bars, with darker bars indicating higher confidence compared to lighter bars (Fig 4). Among 401 amino acids, 63 (15.71%) formed strand, 112 (27.93%) involved in helix formation, and the rest (226, 56.36%) represented coil structures.

**Fig 3.**
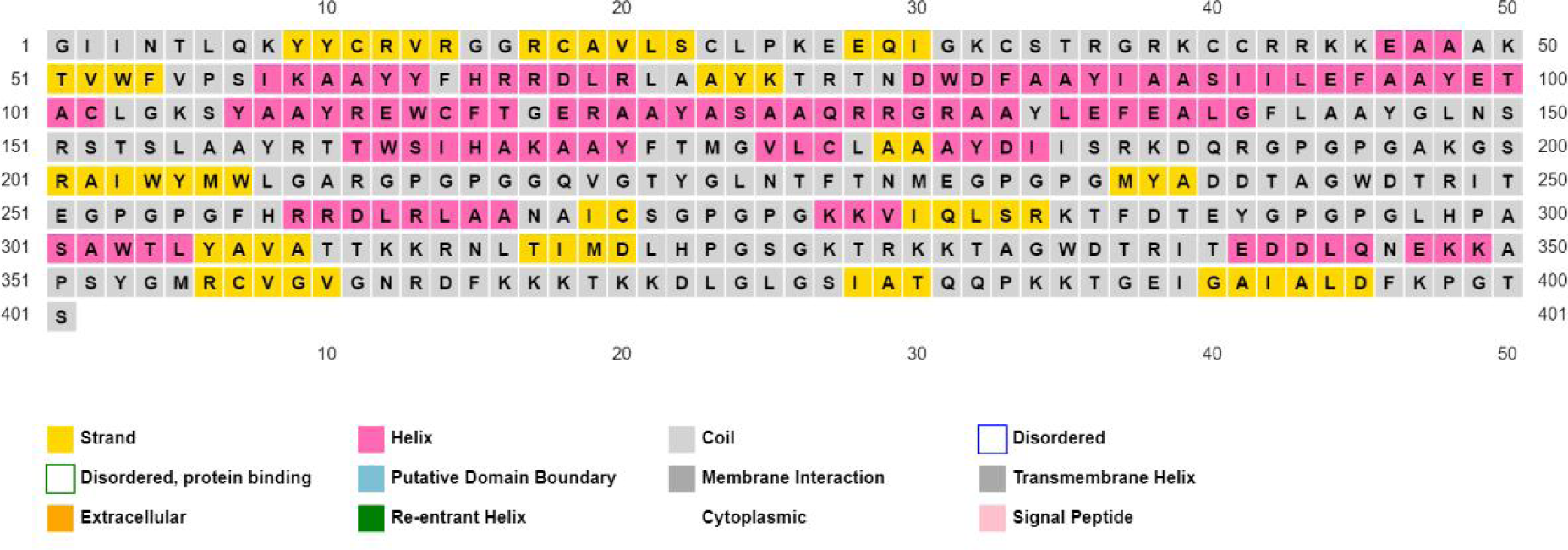
Predicted secondary structure. Different portions are colored based on the predicted structure.

**Fig 4.**
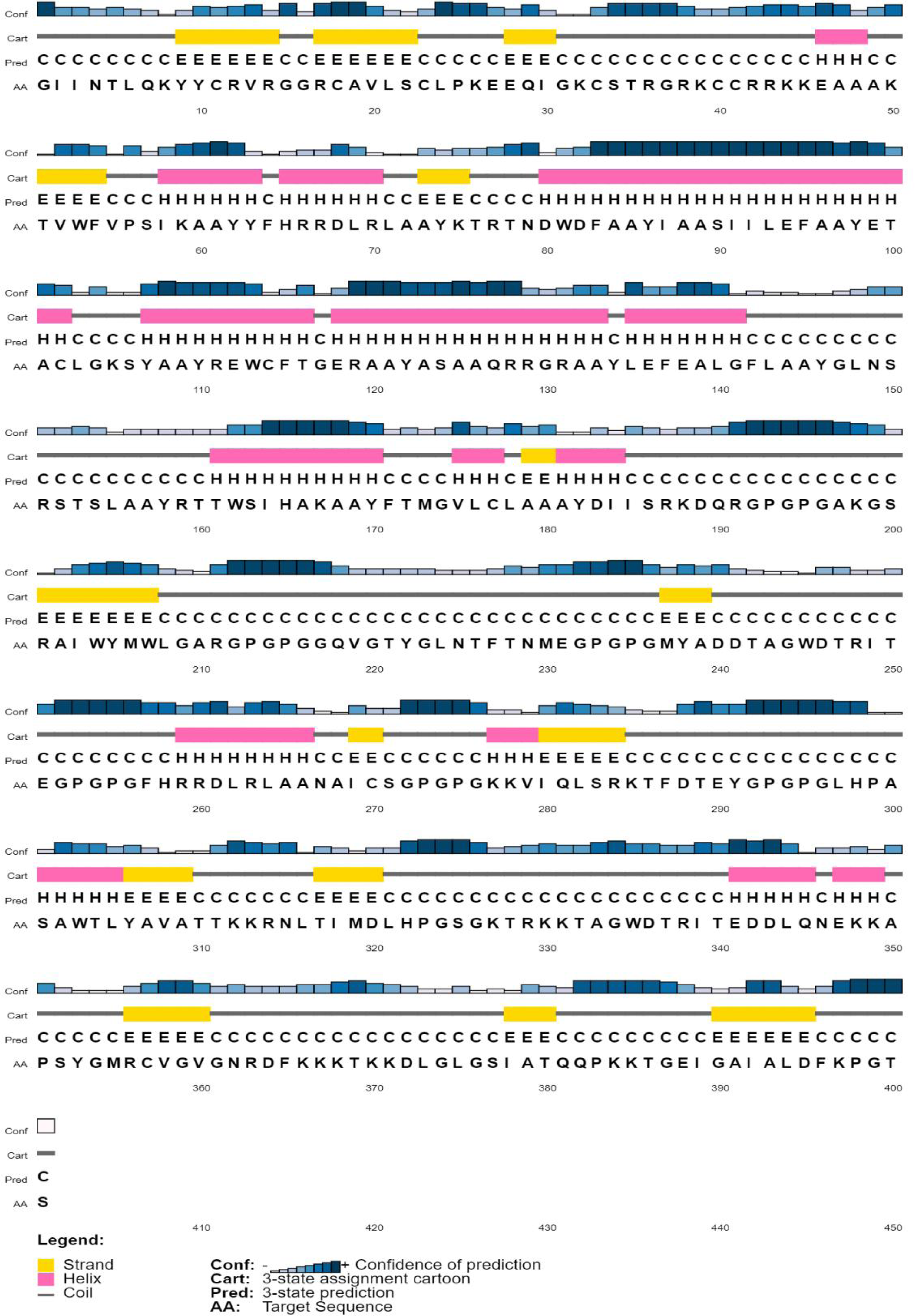
Predicted and secondary structure. Different portions are colored based on the predicted structure.

### Tertiary structure modeling and refinement

The tertiary structure of the protein in PDB format was downloaded from the 3Dpro tool of SCRATCH Protein Predictor. Upon prediction, the structure was refined using the GalaxyRefine tool, which yielded five polished models. Among the refined models, model 4 was selected due to its favorable Rama score of 95.6. Additionally, the GDT-HA score for model 4 was determined to be 0.8953. Moreover, the model has a root mean square deviation (RMSD) of 0.559 and a MolProbity of 2.011. The refined tertiary structure of the vaccine is shown in Fig 5A.

**Fig 5.**
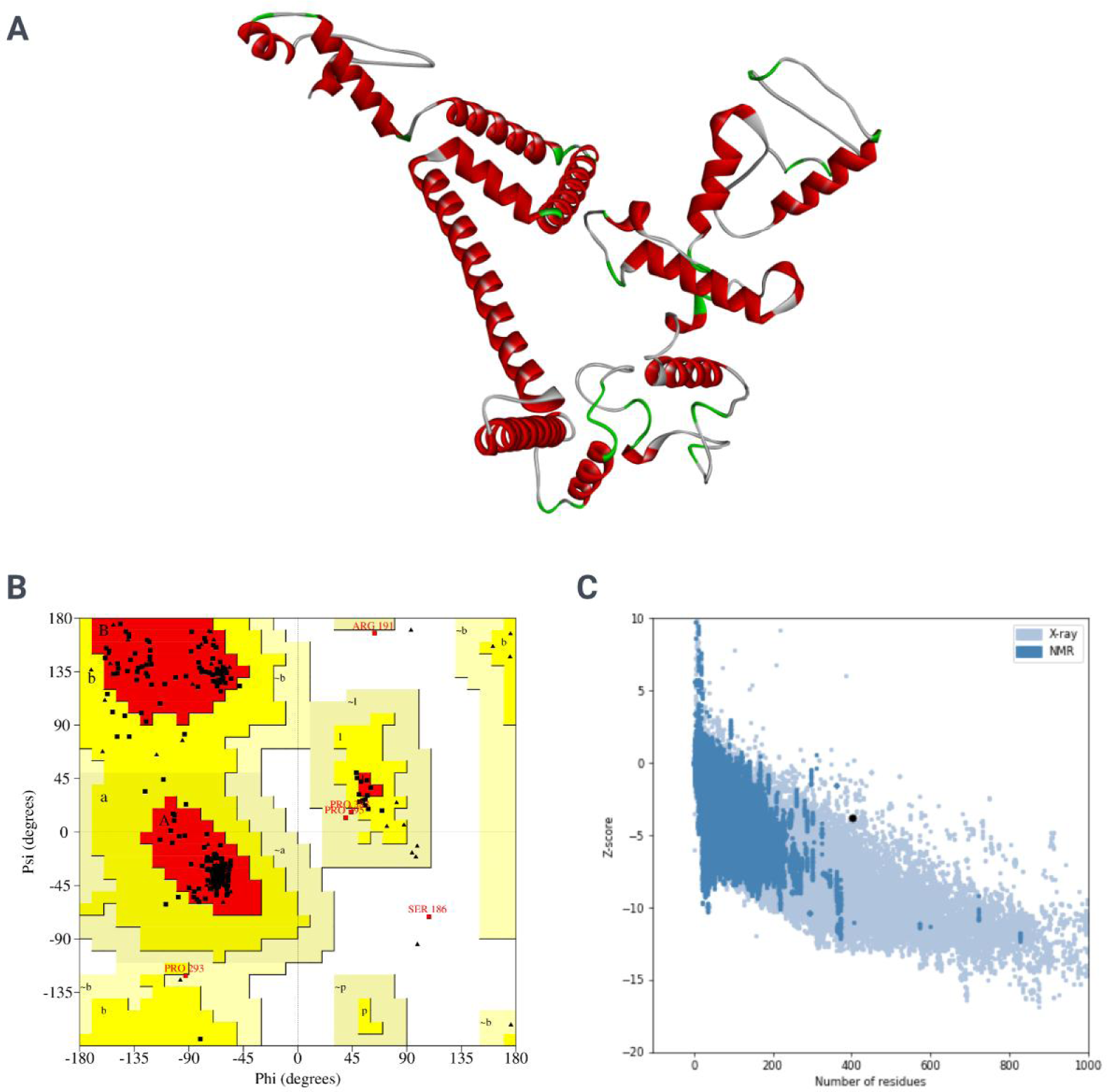
Prediction and validation of tertiary structure. (A) Tertiary refined structure. (B) Ramachandran plots. (C) Z-score plot.

### Tertiary structure validation

The PROCHECK v.3.5 tool of SAVES v6.0 server analyzed the refined structure and illustrated a Ramachandran plot (Fig 5B). The Ramachandran plot depicts that 93.1% (312) of residues were found in the most favored region and 6.3% (21) occupied the additionally allowed region among the considerable 335 (of 401) residues. Notably, no residues were observed in the generously allowed regions and only 0.6% (2) residues were in the disallowed regions, indicating the structural integrity of the peptide. Subsequently, the vaccine construct yielded a Z-score of −3.83 which was retrieved from ProSA-web server (Fig 5C).

### Molecular docking analysis

Once the molecular docking analyses were performed using the ClusPro 2.0 server’s supercomputers to investigate the interactions between the vaccine model and toll-like receptor 2 (TLR2) and TLR4, the vaccine demonstrated significant interactions with both the immune receptors. A total of 30 docked conformations were yielded in the docking operations. The top results had been considered for each operation. The complex formed with TLR2 had a lowest energy of −1393.3 kJ/mol with 76 members. In contrast, the complex formed with TLR4 showed a lowest energy of −1240.5 kJ/mol with 40 members. The docked complexes with TLR2 and TLR4 have been shown in Figs 6 and 7. There were observed 22 residues from the vaccine and 24 residues from TLR2 participating in different types of interactions (Fig 6B and 6C). Specifically, 5 salt bridges, 19 non-bonded contacts, and 167 hydrogen bonds were observed in the vaccine-TLR2 complex (Fig 6C). In contrast, in the vaccine-TLR4 complex, 44 residues from the vaccine and 32 residues from TLR2 were speculated to interact (Fig 7B and 7C). In this complex, 9 salt bridges, 31 non-bonded contacts, and 297 hydrogen bonds were formed (Fig 7C).

**Fig 6.**
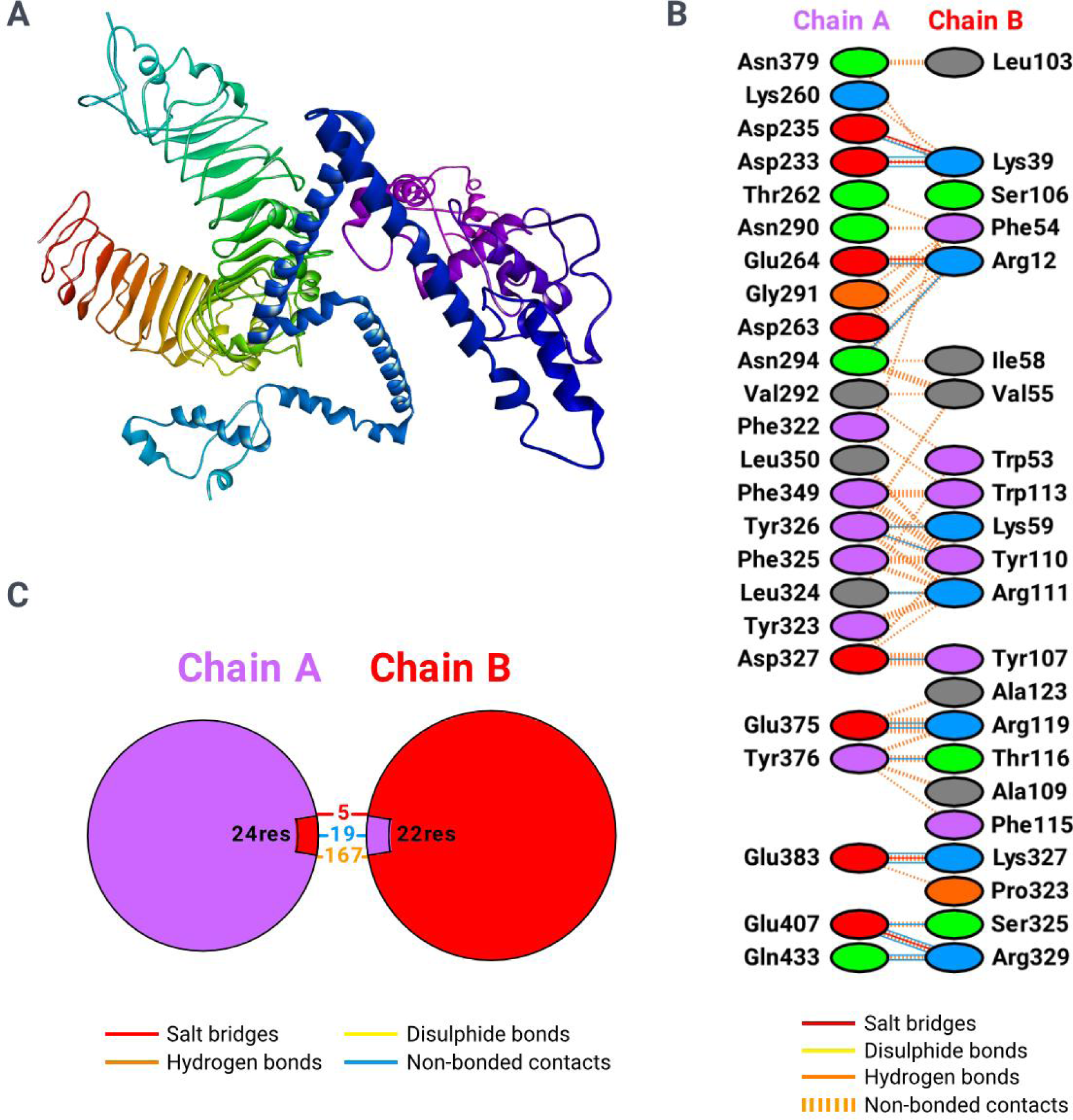
Molecular docking of the vaccine (chain B) with TLR2 (chain A). (A) The three dimensional conformation of vaccine-TLR2 complex. (B) Interactions between the vaccine and TLR2 characterized by types and amino acids. (C) Overall total number of interactions between the vaccine and TLR2 by types.

**Fig 7.**
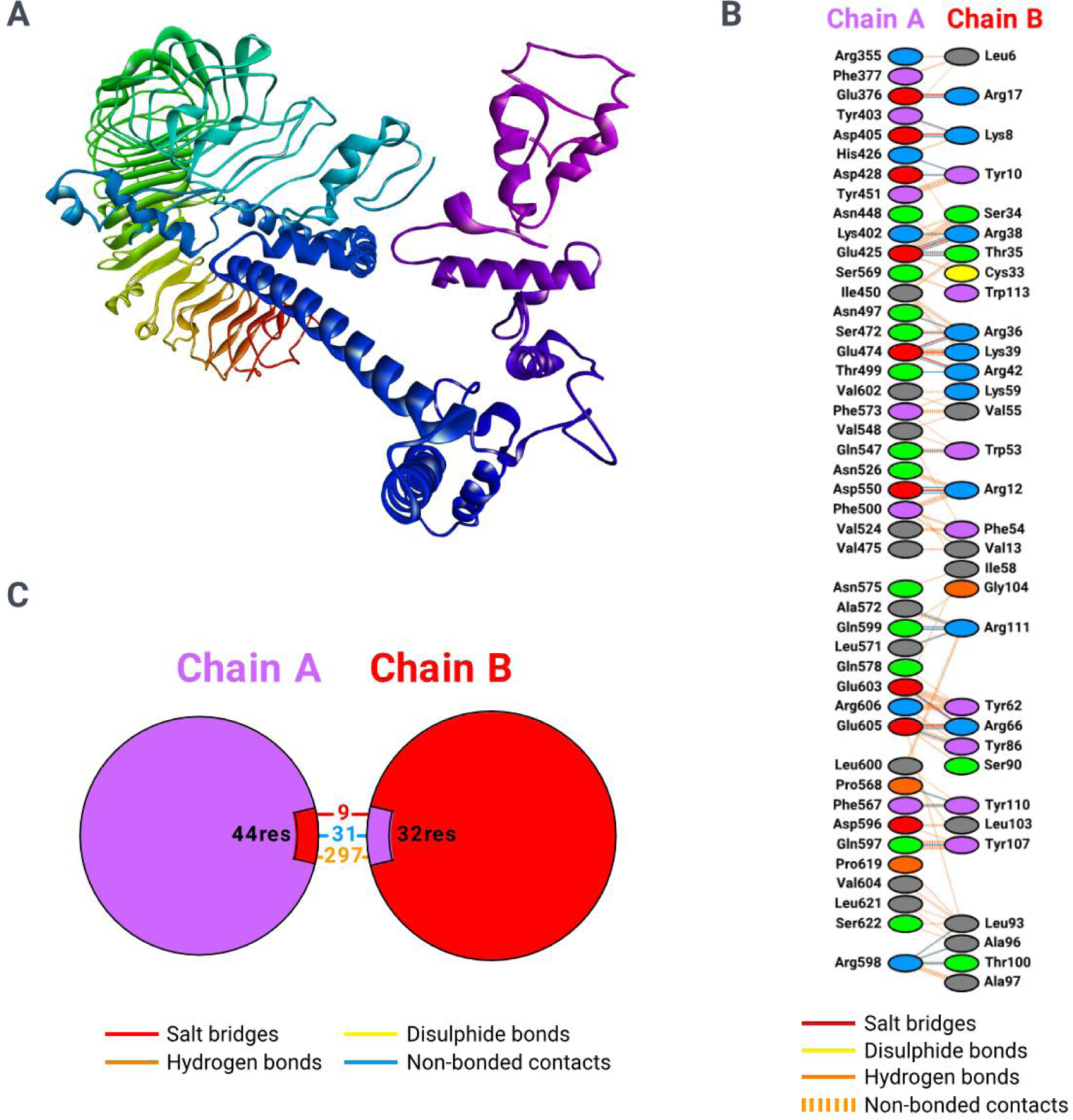
Molecular docking of the vaccine (chain B) with TLR4 (chain A). (A) The three dimensional conformation of vaccine-TLR4 complex. (B) Interactions between the vaccine and TLR4 characterized by types and amino acids. (C) Overall total number of interactions between the vaccine and TLR4 by types.

### Molecular dynamics (MD) simulation analysis

XmGrace package was used to analyze the MD trajectory parameters [68]. During the MD run, the RMSD is calculated between a defined starting point of the simulation and all succeeding frames [69]. The analysis revealed an average RMSD value of 1.187 nm for the vaccine-TLR2 complex and 1.913 nm for the vaccine-TLR4 complex (Figs 8A and 9A). From the graph, the stability of the RMSD value for the vaccine-TLR4 complex can be observed at 25 to 31 nanoseconds (Fig 9A), and for the vaccine-TLR2 complex, it was from 16 to 22 nanoseconds (Fig 8A). The RMSD value is higher in the vaccine-TLR4 complex (1.913 nm) and lower in the vaccine-TLR2 complex (1.187 nm). The RMSD is useful in analyzing the time-dependent motion of a given structure throughout the simulation [70]. In this way, a plateau of RMSD values indicates that the form fluctuates around a stable average conformation, which can be observed in all of the performed MD simulations.

**Fig 8.**
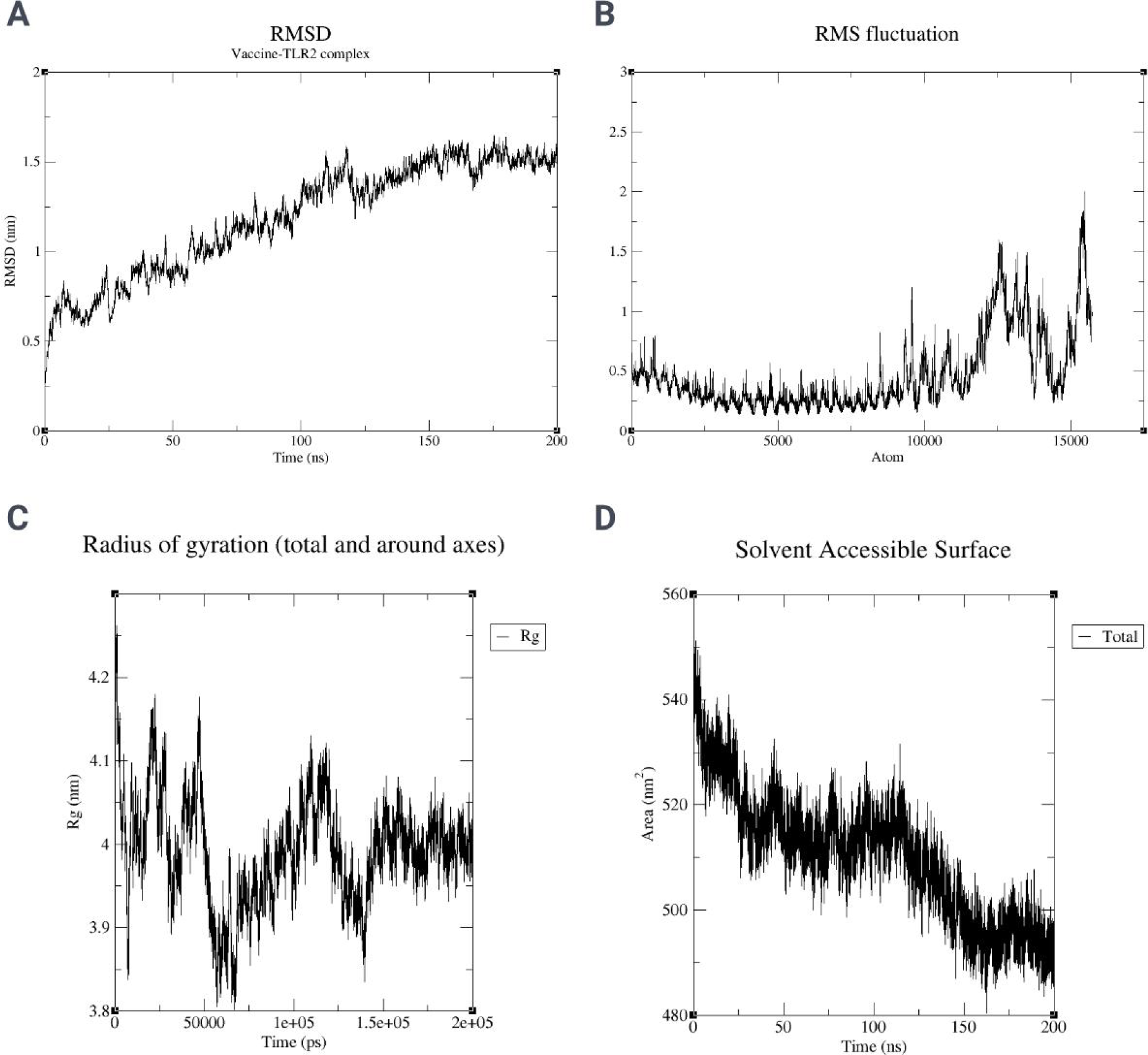
Molecular dynamics simulation graphs of the vaccine with TLR2. (A) Root mean square deviations (RMSD). (B) Root mean square fluctuations (RMSF). (C) Radii of gyration (Rg). (D) Solvent accessible surface area (SASA).

**Fig 9.**
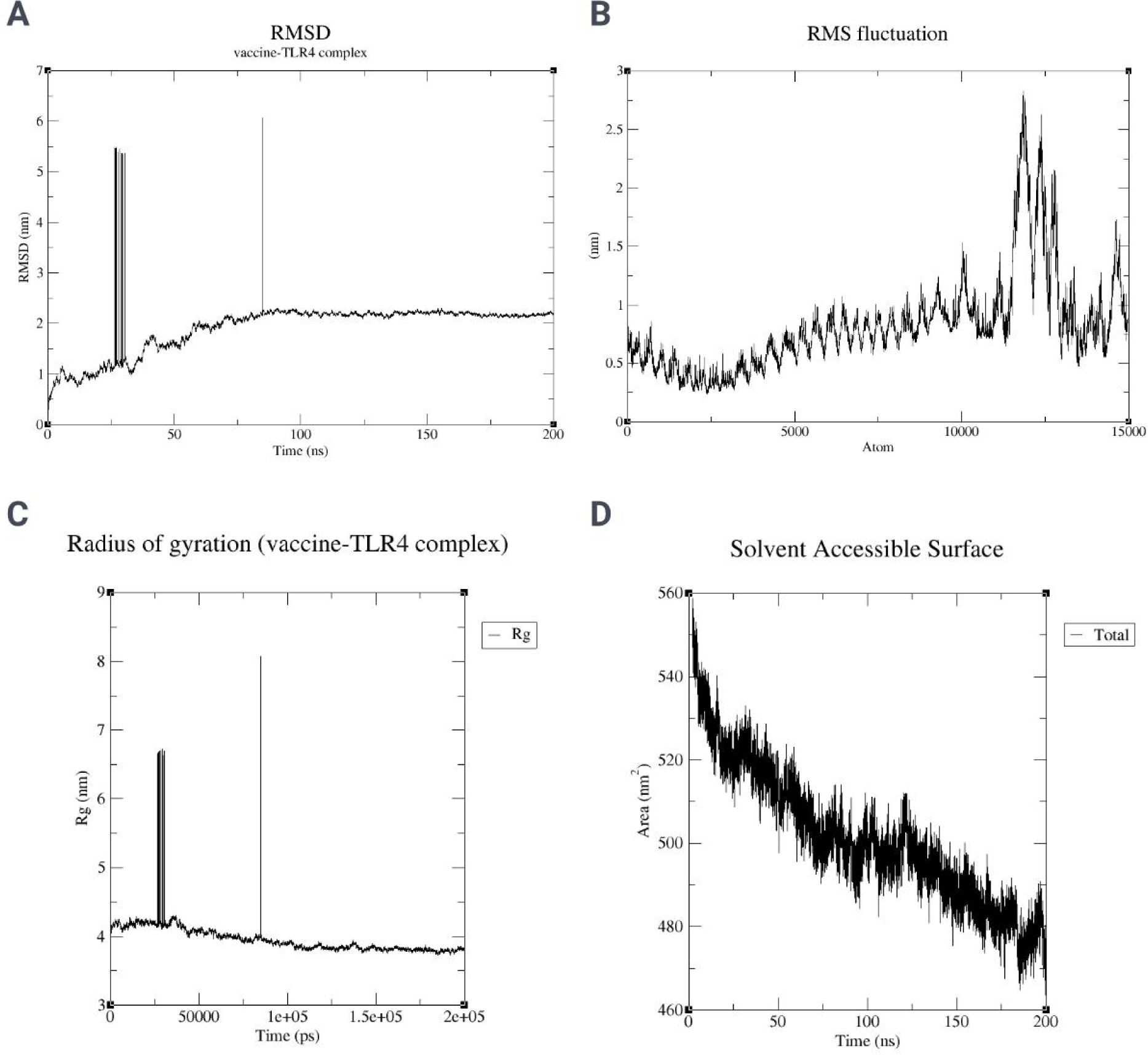
Molecular dynamics simulation graphs of the vaccine with TLR4. (A) Root mean square deviations (RMSD). (B) Root mean square fluctuations (RMSF). (C) Radii of gyration (Rg). (D) Solvent accessible surface area (SASA).

The local flexibility based on residue displacements during the MD simulation can be defined using RMSF values [68]. According to the analysis, the stable RMSF value for the vaccine-TLR2 complex can be seen from the residue range of 5200–8000 (Fig 8B), and for the vaccine-TLR4 complex, the residue range is 6800–7000 (Fig 9B). The average RMSF value for the vaccine-TLR4 complex was 0.640 nm and for the vaccine-TLR2 complex, it was 0.301 nm. Therefore, the RMSF results are higher for the vaccine-TLR4 complex (0.640 nm) than that for the vaccine-TLR2 complex (0.301 nm). Higher RMSF values represent more flexible movements, whereas lower RMSF values represent movements that are more constrained in regards to average locations during simulation [70].

The radius of gyration (RG) is commonly described as the root mean square distance of a group of atoms from their shared center of mass, with the added consideration of mass weighting [71]. According to the results of the RG analysis, the stable RG value for the vaccine-TLR4 complex is from 115 to 133 nanoseconds (Fig 9C), and for the vaccine-TLR2 complex, it is from 150 to 165 nanoseconds (Fig 8C). the average RG value for the vaccine-TLR4 complex was 3.957 nm and for the vaccine-TLR2 complex was 3.99 nm. As a result, the RG score was higher in the vaccine-TLR2 complex (3.99 nm) compared to the vaccine-TLR4 complex (3.957 nm). The findings suggest that the vaccine-TLR4 complex exhibited lower compactness compared to the apoprotein.

The SASA, or solvent-accessible surface area, is widely recognized as the portion of a given protein that is accessible to the surrounding solvent. The SASA analysis methodology offers insights into the ability of a protein to engage in molecular interactions [71]. The results of the SASA analysis indicate that the stable SASA value range for the vaccine-TLR4 complex was 75 to 130 nanoseconds and for the vaccine-TLR2 complex, it was 50 to 110 nanoseconds. The average SASA of the vaccine-TLR2 complex was 510.15 nm^2^ (Fig 8D), and for the vaccine-TLR4 complex was 501.11 nm^2^ (Fig 9D). Hence, the SASA score was more significant for the vaccine-TLR2 complex (510.15 nm^2^) than that of the vaccine-TLR4 complex (501.11 nm^2^). The findings indicate that the vaccine-TLR4 complex exhibit comparatively lower accessibility in comparison to the vaccine-TLR2 complex, potentially impacting their ability to engage with other molecules.

### Immune simulation analysis

The graphs of immune simulations depicted in Figs 8A–F and 9A–F were retrieved from the C-IMMSIM online server. Upon three consecutive doses at 20-day intervals, the vaccine was observed to initiate a host immune response to a good extent. The B-cell population per cubic millimeter was seen to elicit the total cell count at the time of injection, which was observed to remain within a consistent range above 400/mm^3^ (Fig 10A). The PLB cell population also evoked a constant value for a long period (Fig 10B). The per-state active B-cell count was detected to remain consistently within 400–500/mm^3^ after 100 days of injection (Fig 10C). The total number of helper T-cells retained a fixed range near 1400/mm^3^ after 4 months of the first injection (Fig 10D). However, after the same period, the non-memory helper T-cells were seen to gradually increase in number, whereas the memory cells were observed to abate. As per the per-state helper T-cell population graph (Fig 10E), the active T-cell population after 4 months was seen to lessen slowly compared to the rapid fall during the previous 2 months. In contrast, the resting cells were observed to increase. Moreover, the per-state active and resting regulatory T-cell population was noticed to uplift near 20/mm^3^ after 6 months from the first dose of vaccination (Fig 10F).

**Fig 10.**
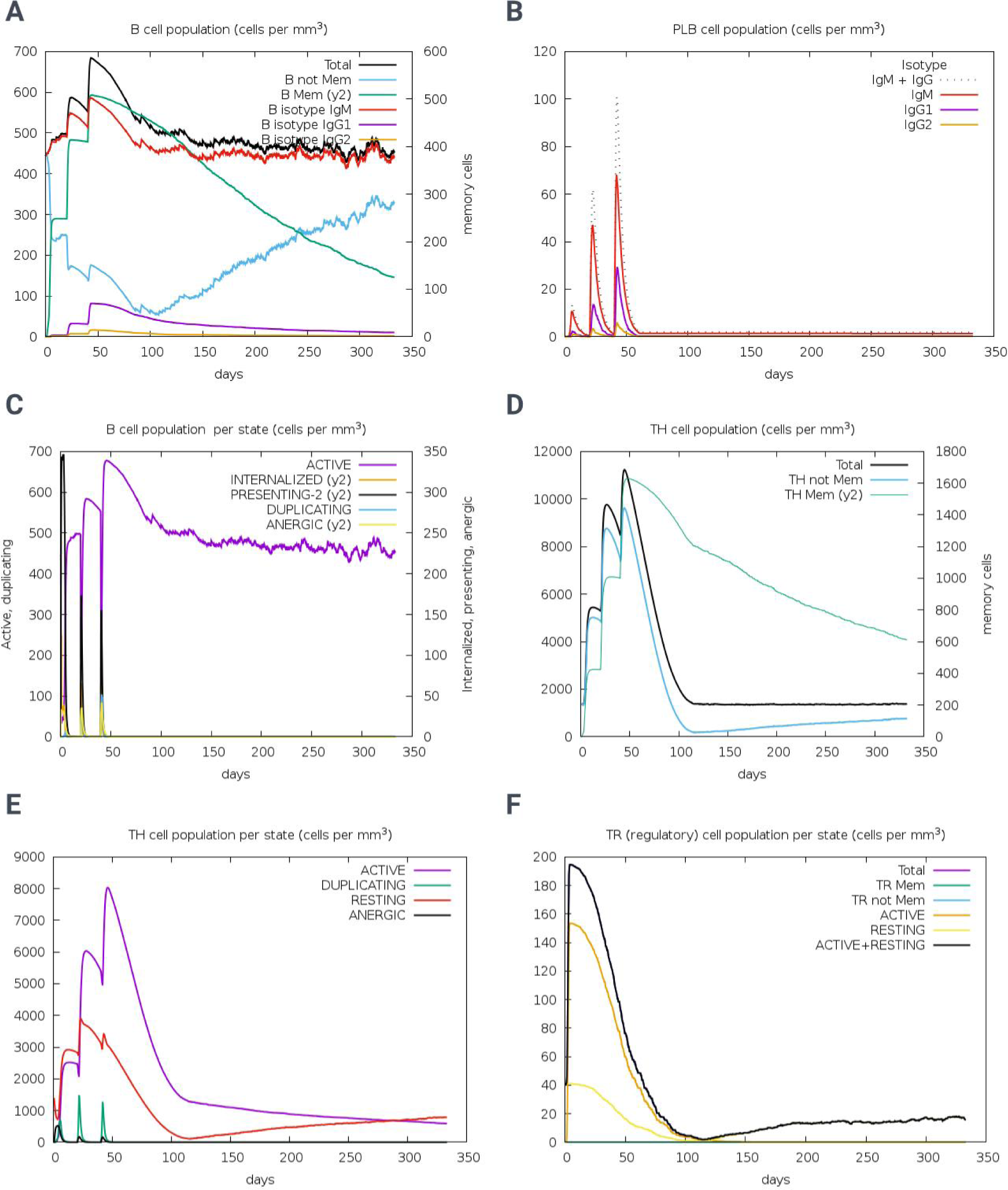
Immune simulation graphs of the vaccine.

The cytotoxic T-cell population fluctuated within 1055–1155/mm^3^, peaking near the 13^th^ day from the first dose (Fig 11A). The balance between active and resting cytotoxic T-cells has been observed in Fig 11B. The natural killer (NK) cell population oscillates within 309– 381/mm^3^ (Fig 11C). The per-state MA and DC populations were observed to elicit an immune response with a significant number of cells (Fig 11D and 11E). Furthermore, the immunoglobulin M (IgM) and IgG were also found to initiate a sufficient immune response for a prolonged period (Fig 11F). Throughout the simulation, the total number of IgM and IgG remained consistently above 10,000/mL These findings validate the immune response-generating efficacy of the designed vaccine candidate.

**Fig 11.**
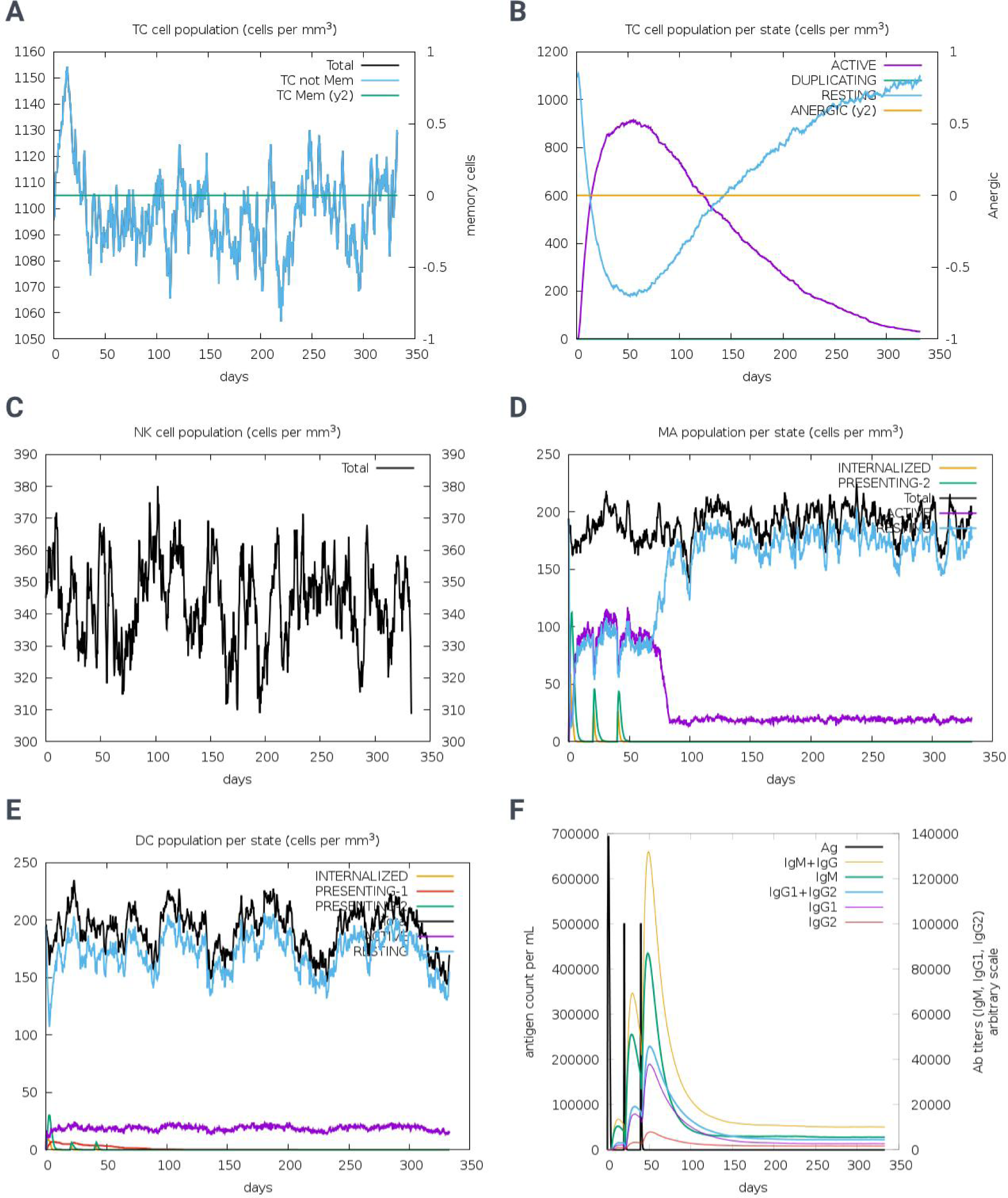
Immune simulation graphs of the vaccine.

### Back-translation, codon optimization, and *in silico* cloning

EMBOSS Backtranseq provided a back-translated RNA sequence of the vaccine with *Escherichia coli* (strain K12) organism. For *in silico* cloning, *E. coli* expression system plays vital role in expressing vaccines efficiently. The back-translated sequence is composed of 1203 nucleotides against the 401 amino acids of the vaccine (<u>S2 File</u>). Afterwards, JCAT was employed for the codon usage adaptation. With *E. coli* (strain K12) expression system, the codon adaptation index (CAI) of the improved sequence was observed as 1.0 and the GC content was 53.78%, whereas the optimum range for CAI is 0.8–1.0 and that of GC content is 30–70%. Before improvement, the CAI and GC content were 0.58 and 60.78%, respectively. These measures signify *E. coli* expression system as efficient and good fit for expressing the designed vaccine. Two cut sites for restriction enzymes *Nde*I and *Xba*I were integrated to the terminals of the vaccine and the DNA turned into 1215 bp in length (Fig 12A). Thus, the restriction enzymes *Nde*I and *Xba*I were common in both the vaccine DNA and the vector. Prior to insertion, the positions of these two enzyme sites in the vector were 238 and 335, respectively, whereas the vector was 5369 bp in length. Upon cutting the vaccine DNA and the vector with the same enzymes, the vaccine DNA was successfully inserted into the pET-28a(+) vector using SnapGene software. The final plasmid length turned into 6480 bp (Fig 12B).

**Fig 12.**
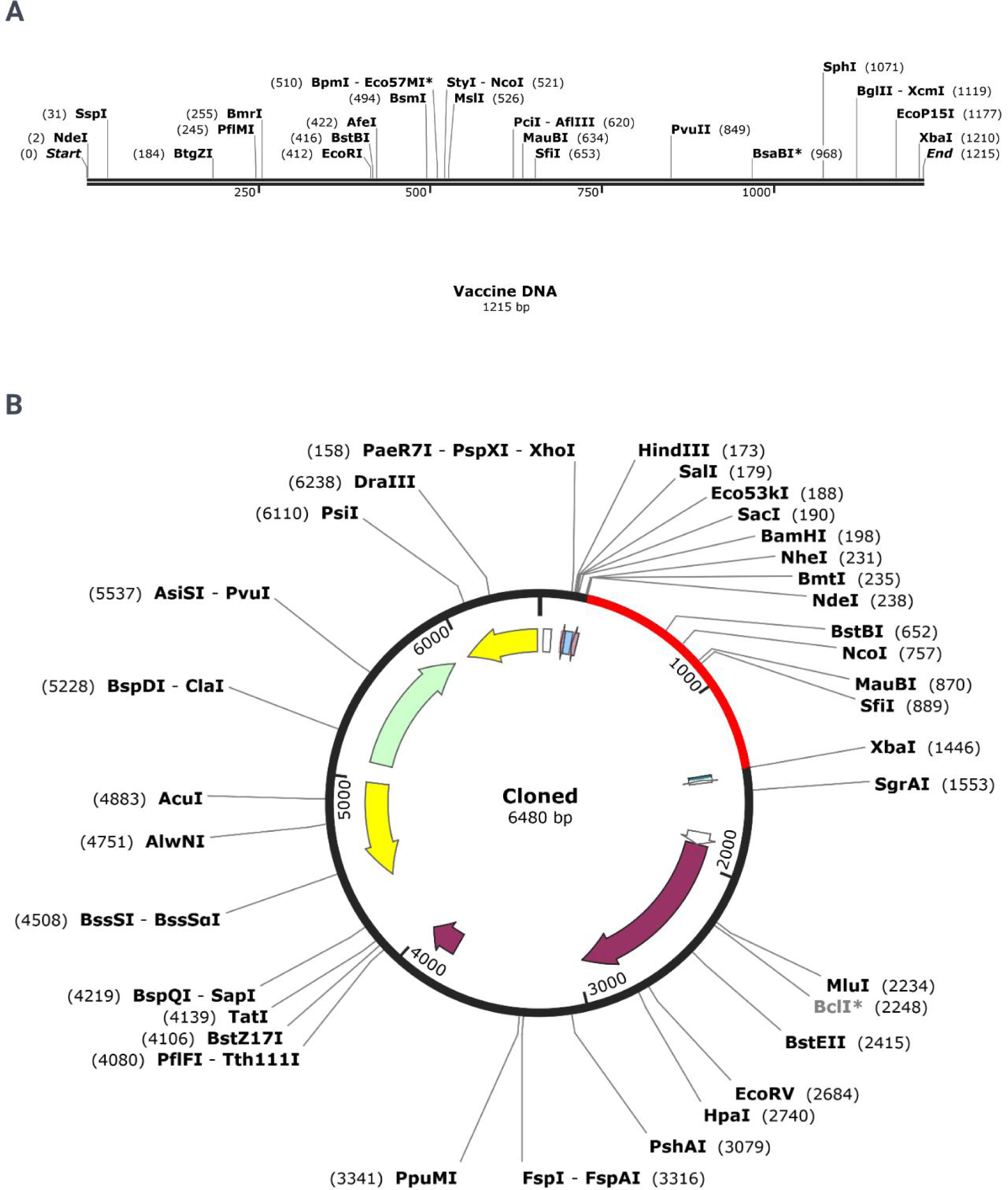
Insertion of the designed vaccine into the pET-28a(+) vector using SnapGene software (www.snapgene.com). (A) The vaccine DNA map after integration of the cut sites. (B) The pET-28a(+) vector map after insertion of the vaccine. The inserted portion iso marked with red color.

## Discussion

Dengue fever can lead to severe flu-like symptoms and even may develop life-threatening dengue hemorrhagic fever or dengue shock syndrome. Every dengue infection carries the risk of hospitalization and severe illness, and if not treated promptly, severe dengue can lead to death [72]. This highlights the importance of vaccination as a preventive measure. The complexity of the DENV, which has four distinct serotypes, poses a significant challenge for vaccine development [73]. Immunity to one serotype does not guarantee protection against the others, and subsequent infections with different serotypes can increase the risk of severe dengue. This necessitates a tetravalent vaccine that can confer immunity against all four serotypes to effectively reduce the global health burden of dengue. Computational approaches have revolutionized vaccine development, offering a faster and more cost-effective route compared to traditional methods [74]. Immunoinformatics and computational vaccinology enable researchers to analyze the genetic makeup of viruses and predict which viral components are most likely to produce an immune response. These methods facilitate the identification of potential B-cell and T-cell epitopes, which are crucial for the design of epitope-based vaccines. Moreover, computational tools allow for the simulation of vaccine behavior in the human body, predicting how the immune system will respond to a vaccine candidate before it is produced and tested in the laboratory or clinical trials. This can significantly accelerate the vaccine development process, making it possible to respond more quickly to emerging infectious diseases.

The proposed vaccine construct, comprising 401 amino acids, includes a combination of 23 epitopes—18 T-cell and 5 B-cell epitopes—thus ensuring broad coverage against all four DENV serotypes. The inclusion of human β-defensin 3 as an adjuvant is a novel approach that could enhance the vaccine’s immunogenicity [75]. The physicochemical properties and solubility of a vaccine construct are pivotal in determining its suitability for clinical use. The vaccine construct in question, with a molecular weight of 43.84 kDa exhibits a robust stability profile, as indicated by an instability index of 28.06 and a GRAVY value of −0.444, suggesting a hydrophilic nature. The solubility scores from SOLpro, SoluProt 1.0, and Protein-Sol further corroborate its soluble characteristics, which is essential for effective delivery and distribution within the body [76].

Antigenicity is a critical factor for vaccine efficacy, and with a score of 0.9319 from VaxiJen 2.0, the vaccine surpasses the threshold for potential immunogenicity [27]. The allergenicity assessments from AllerTOP v.2 and AlgPred server classify the vaccine as a probable non-allergen, which is crucial for minimizing adverse reactions. The absence of toxicity, as indicated by the lack of results from the T3DB server, further supports the vaccine’s safety profile. Population coverage is a measure of a vaccine’s potential reach and impact. With a reported 97.35% worldwide population coverage, the vaccine demonstrates a promising capacity for widespread immunization, which is essential for controlling the spread of the targeted pathogen3. The average hits and PC90 values for both MHC-I and MHC-II epitopes suggest a broad recognition by the human leukocyte antigen (HLA) system, which is fundamental for eliciting a robust immune response3. Disulfide bonds contribute to the structural stability of proteins, and their presence in a vaccine construct can enhance its stability and efficacy. The prediction of 4 disulfide bonds within the vaccine sequence, involving cysteines at specific positions, suggests a well-defined tertiary structure conducive to proper folding and function [77, 78]. This structural integrity is vital for the vaccine’s ability to induce an appropriate immune response.

The secondary structure prediction for the vaccine peptide, as generated by the PSIPRED 4.0 server, provides a foundational understanding of the protein’s conformational layout. The distribution of helices, strands, and coils within the peptide is crucial for its function, with the confidence levels indicated by the color bars offering insights into the reliability of these predictions [79]. The tertiary structure modeling and refinement process, utilizing the 3Dpro tool and GalaxyRefine, has resulted in a polished structure with a favorable Rama score and GDT-HA score, indicating a high-quality model suitable for further analysis. The root mean square deviation (RMSD) and MolProbity score further support the structural integrity of the refined model. Validation of the tertiary structure through PROCHECK and the Ramachandran plot analysis reveals a high percentage of residues in the most favored regions, which is indicative of a well-folded protein. The Z-score obtained from the ProSA-web server aligns with the expected range for similar-sized proteins, confirming the model’s validity.

The molecular docking analysis, performed using ClusPro 2.0, has demonstrated significant interactions between the vaccine model and toll-like receptors TLR2 and TLR4. The binding energies and the number of members in the docked conformations suggest a stable and potentially effective interaction, which is essential for the vaccine’s ability to elicit an immune response. Molecular dynamics (MD) simulations provide critical insights into the dynamic behavior and stability of vaccine-receptor complexes, which are essential for understanding their potential efficacy. The analysis of the vaccine-TLR2 and vaccine-TLR4 complexes reveals distinct stability and flexibility profiles that could influence their immunogenicity and interaction with the immune system. The RMSD values for the vaccine-TLR2 (average of 1.187 nm) and vaccine-TLR4 (average of 1.913 nm) complexes suggest that the vaccine-TLR2 complex maintains a more stable conformation throughout the simulation period. The observed plateau in RMSD values indicates that both complexes achieve a stable average conformation after initial fluctuations. The RMSF analysis indicates that the vaccine-TLR4 complex (average of 0.640 nm) exhibits greater local flexibility compared to the vaccine-TLR2 complex (average of 0.301 nm). This flexibility could be advantageous for the vaccine-TLR4 complex, potentially allowing for better accommodation of epitope variations and enhancing immune recognition. The RG values suggest that the vaccine-TLR2 complex (average of 3.99 nm) is slightly more compact than the vaccine-TLR4 complex (average of 3.957 nm). A higher RG value typically correlates with a less compact structure, which may affect the stability and immunogenicity of the complex. The SASA analysis shows that the vaccine-TLR2 complex has a higher average SASA (510.15 nm²) compared to the vaccine-TLR4 complex (501.11 nm²). A larger SASA may indicate a higher degree of exposure to solvent molecules, which could facilitate interactions with immune cells and potentially enhance the immunogenic response.

The immune simulation analysis, as depicted in Figures 10 and 11, provides a quantitative assessment of the host immune response following administration of a novel vaccine candidate. The data, derived from the C-IMMSIM online server, underscores the vaccine’s ability to elicit a robust and sustained immune response, characterized by significant B-cell and T-cell activation over an extended period. The B-cell response, with populations remaining above 400/mm^3^, indicates a strong initial immune reaction, which is crucial for the formation of memory cells and long-term immunity. The persistence of active B-cells within 400–500/mm^3^ after 100 days suggests the potential for lasting antibody-mediated immunity, which is essential for protection against reinfection. Helper T-cells, maintaining a steady range near 1400/mm^3^, reflect a consistent cell-mediated immune response, which is vital for the activation of other immune cells. The increase in non-memory helper T-cells and the decline in memory cells may indicate a shift towards an active response phase, possibly due to ongoing antigenic stimulation. Regulatory T-cells, increasing to about 20/mm^3^ after 6 months, suggest a mechanism to modulate the immune response, preventing potential autoimmunity or hyperactivation. The fluctuation of cytotoxic T-cells within 1055–1155/mm^3^, peaking around the 13th day, demonstrates their role in the direct elimination of infected cells. The oscillation of NK cells within 309–381/mm^3^ highlights their importance in the early stages of the immune response, providing rapid action against virally infected cells. The consistent levels of IgM and IgG above 10,000/mL throughout the simulation period indicate a strong humoral response, which is critical for neutralizing pathogens and preventing their spread.

### Conclusion

This study has resulted in the successful development of a tetravalent multi-epitope subunit vaccine for all DENV serotypes. Notably, the vaccine has an antigenicity score of 0.9319 which is the highest among all computationally developed vaccines to date. Moreover, with a worldwide population coverage of 97.35%, the vaccine tends to be an efficacious candidate for any ethnicity. The physicochemical properties and solubility of the vaccine were observed in the optimum ranges. The molecular docking results and MD simulations showed propitious interactions, stability, and flexibility with immune receptors (TLR2 and TLR4) of the human body. Furthermore, immune simulations showed that the vaccine is capable of eliciting proper immune responses in the human body for a long time span. Finally, the designed vaccine was successfully cloned by inserting it into the pET-28a(+) vector of *E. coli* (strain K12). Importantly, this vaccine candidate shows the most propitious results among all DENV vaccines developed till today. However, the data explored, assessed and the simulations ran in this study were parts of reverse vaccinology and immunoinformatics approach. Thus, in vitro and in vivo experimental validations are required to confirm the efficacy, immunogenicity, and toxicity of the designed vaccine. This study will work as the basis of future validations, that might bring a scientific breakthrough.

## Data availability

The complete genome sequences of all four serotypes of the virus that are retrieved from NCBI website (https://www.ncbi.nlm.nih.gov/) are included in <u>S1 File</u>. Required validation data are included in <u>S2 File</u> and <u>S1 Fig</u>. Additional data are available on requests.

## Acknowledgments

The authors are thankful to Professor Dr. Sukalyan Kumar Kundu for his keen supervision and providing the facilities to conduct the research successfully.

## Author contributions

**Conceptualization:** Sukalyan Kumar Kundu, Tanbin Jahan Ferdousy, Md Mehedy Hasan Miraz

**Data Curation:** Tanbin Jahan Ferdousy, Md Afif Ullah, Md Tahsinul Haque Risat, Shouhardyo Kundu

**Formal Analysis:** Md Mehedy Hasan Miraz, Tawsif Al Arian, Md Afif Ullah

**Investigation:** Tanbin Jahan Ferdousy

**Methodology:** Md Mehedy Hasan Miraz, Tawsif Al Arian, Md Tahsinul Haque Risat, Md Afif Ullah

**Project Administration:** Sukalyan Kumar Kundu, Tanbin Jahan Ferdousy, Abul Bashar Ripon Khalipha

**Resources:** Tanbin Jahan Ferdousy, Md Mehedy Hasan Miraz, Tawsif Al Arian, Most. Afrin Akter, Md Tahsinul Haque Risat

**Software:** Md Mehedy Hasan Miraz, Tanbin Jahan Ferdousy, Tawsif Al Arian, Most. Afrin Akter

**Supervision:** Sukalyan Kumar Kundu

**Validation:** Tanbin Jahan Ferdousy, Bidduth Kumar Sarkar, Arghya Prosun Sarkar, Abul Bashar Ripon Khalipha

**Visualization:** Tanbin Jahan Ferdousy, Md Mehedy Hasan Miraz, Shouhardyo Kundu **Writing – Original Draft Preparation:** Tanbin Jahan Ferdousy, Md Mehedy Hasan Miraz **Writing – Review & Editing:** Sukalyan Kumar Kundu, Tanbin Jahan Ferdousy

## Supporting information captions

**S1 File. Complete genome sequences of DENV-1 to −4.**

**S2 File. Back-translated and codon-optimized sequence of the vaccine.**

**S1 Fig. Population coverage graphs retrieved from IEDB Population server.**

## References

1. CDC. Data and Maps | Dengue | CDC [Internet]. 2024 [cited 2024 Mar 24]. Available from: https://www.cdc.gov/dengue/statistics-maps/data-and-maps.html

2. WHO. Dengue - Global situation [Internet]. 2023 Dec. Available from: https://www.who.int/emergencies/disease-outbreak-news/item/2023-DON498

3. Guzman MG, Gubler DJ, Izquierdo A, Martinez E, Halstead SB. Dengue infection. Nat Rev Dis Primers. 2016 Aug 18;2(1):16055.

4. WHO. Dengue and severe dengue [Internet]. 2023 Mar [cited 2024 Mar 25]. Available from: https://www.who.int/news-room/fact-sheets/detail/dengue-and-severe-dengue

5. Byk LA, Gamarnik AV. Properties and Functions of the Dengue Virus Capsid Protein. Annu Rev Virol. 2016 Sep 29;3(1):263–81.

6. Dias RS, Teixeira MD, Xisto MF, Prates JWO, Silva JDD, Mello IO, et al. DENV-3 precursor membrane (prM) glycoprotein enhances E protein immunogenicity and confers protection against DENV-2 infections in a murine model. Human Vaccines & Immunotherapeutics. 2021 May 4;17(5):1271–7.

7. Young E, Yount B, Pantoja P, Henein S, Meganck RM, McBride J, et al. A live dengue virus vaccine carrying a chimeric envelope glycoprotein elicits dual DENV2-DENV4 serotype-specific immunity. Nat Commun. 2023 Mar 13;14(1):1371.

8. Alcalá AC, Maravillas JL, Meza D, Ramirez OT, Ludert JE, Palomares LA. Dengue Virus NS1 Uses Scavenger Receptor B1 as a Cell Receptor in Cultured Cells. Heise MT, editor. J Virol. 2022 Mar 9;96(5):e01664-21.

9. Xie X, Zou J, Zhang X, Zhou Y, Routh AL, Kang C, et al. Dengue NS2A Protein Orchestrates Virus Assembly. Cell Host & Microbe. 2019 Nov;26(5):606–622.e8.

10. Li Q, Kang C. Structures and Dynamics of Dengue Virus Nonstructural Membrane Proteins. Membranes. 2022 Feb 17;12(2):231.

11. Swarbrick CMD, Basavannacharya C, Chan KWK, Chan SA, Singh D, Wei N, et al. NS3 helicase from dengue virus specifically recognizes viral RNA sequence to ensure optimal replication. Nucleic Acids Research. 2017 Dec 15;45(22):12904–20.

12. Cortese M, Mulder K, Chatel-Chaix L, Scaturro P, Cerikan B, Plaszczyca A, et al. Determinants in Nonstructural Protein 4A of Dengue Virus Required for RNA Replication and Replication Organelle Biogenesis. López S, editor. J Virol. 2021 Oct 13;95(21):e01310-21.

13. Bhatnagar P, Sreekanth GP, Murali-Krishna K, Chandele A, Sitaraman R. Dengue Virus Non-Structural Protein 5 as a Versatile, Multi-Functional Effector in Host–Pathogen Interactions. Front Cell Infect Microbiol. 2021 Mar 18;11:574067.

14. McArthur MA, Sztein MB, Edelman R. Dengue vaccines: recent developments, ongoing challenges and current candidates. Expert Review of Vaccines. 2013 Aug;12(8):933–53.

15. Freedman DO. A new dengue vaccine (TAK-003) now WHO recommended in endemic areas; what about travellers? Journal of Travel Medicine. 2023 Nov 18;30(7):taad132.

16. Santé WHO= O mondiale de la. Weekly Epidemiological Record, 2023, vol. 98, 47 [full issue]. Vol. 98, Weekly Epidemiological Record = Relevé épidémiologique hebdomadaire. World Health Organization = Organisation mondiale de la Santé; 2023. p. 599–620.

17. Santé WHO= O mondiale de la. Dengue vaccine: WHO position paper – September 2018 – Note de synthèse de l’OMS sur le vaccin contre la dengue– septembre 2018. Vol. 93, Weekly Epidemiological Record = Relevé épidémiologique hebdomadaire. World Health Organization = Organisation mondiale de la Santé; 2018. p. 457–76.

18. Redoni M, Yacoub S, Rivino L, Giacobbe DR, Luzzati R, Di Bella S. Dengue: Status of current and under-development vaccines. Reviews in Medical Virology. 2020 Jul;30(4):e2101.

19. Angelin M, Sjölin J, Kahn F, Ljunghill Hedberg A, Rosdahl A, Skorup P, et al. Qdenga® - A promising dengue fever vaccine; can it be recommended to non-immune travelers? Travel Medicine and Infectious Disease. 2023 Jul;54:102598.

20. Tricou V, Yu D, Reynales H, Biswal S, Saez-Llorens X, Sirivichayakul C, et al. Long-term efficacy and safety of a tetravalent dengue vaccine (TAK-003): 4·5-year results from a phase 3, randomised, double-blind, placebo-controlled trial. The Lancet Global Health. 2024 Feb;12(2):e257–70.

21. Wang Y, Ling L, Zhang Z, Marin-Lopez A. Current Advances in Zika Vaccine Development. Vaccines. 2022 Oct 28;10(11):1816.

22. Malik S, Kishore S, Nag S, Dhasmana A, Preetam S, Mitra O, et al. Ebola Virus Disease Vaccines: Development, Current Perspectives & Challenges. Vaccines. 2023 Jan 26;11(2):268.

23. DeFilippis VR. Chikungunya Virus Vaccines: Platforms, Progress, and Challenges. In Berlin, Heidelberg: Springer Berlin Heidelberg; 2019 [cited 2024 Apr 13]. (Current Topics in Microbiology and Immunology). Available from: https://link.springer.com/10.1007/82_2019_175

24. Islam MdA. A review of SARS-CoV-2 variants and vaccines: Viral properties, mutations, vaccine efficacy, and safety. Infectious Medicine. 2023 Dec;2(4):247–61.

25. Pintado Silva J, Fernandez-Sesma A. Challenges on the development of a dengue vaccine: a comprehensive review of the state of the art. Journal of General Virology [Internet]. 2023 Mar 1 [cited 2024 Apr 13];104(3). Available from: https://www.microbiologyresearch.org/content/journal/jgv/10.1099/jgv.0.001831

26. Benson DA, Cavanaugh M, Clark K, Karsch-Mizrachi I, Lipman DJ, Ostell J, et al. GenBank. Nucleic Acids Research. 2012 Nov 26;41(D1):D36–42.

27. Doytchinova IA, Flower DR. VaxiJen: a server for prediction of protective antigens, tumour antigens and subunit vaccines. BMC Bioinformatics. 2007 Dec;8(1):4.

28. Saha S, Raghava GPS. Prediction of continuous B-cell epitopes in an antigen using recurrent neural network. Proteins. 2006 Oct;65(1):40–8.

29. Dhanda SK, Mahajan S, Paul S, Yan Z, Kim H, Jespersen MC, et al. IEDB-AR: immune epitope database—analysis resource in 2019. Nucleic Acids Research. 2019 Jul 2;47(W1):W502–6.

30. Dimitrov I, Bangov I, Flower DR, Doytchinova I. AllerTOP v.2—a server for in silico prediction of allergens. J Mol Model. 2014 Jun;20(6):2278.

31. Wold S, Jonsson J, Sjörström M, Sandberg M, Rännar S. DNA and peptide sequences and chemical processes multivariately modelled by principal component analysis and partial least-squares projections to latent structures. Analytica Chimica Acta. 1993 May;277(2):239–53.

32. Venkatarajan MS, Braun W. New quantitative descriptors of amino acids based on multidimensional scaling of a large number of physical–chemical properties. Molecular modeling annual. 2001 Dec 1;7(12):445–53.

33. Bui HH, Sidney J, Li W, Fusseder N, Sette A. Development of an epitope conservancy analysis tool to facilitate the design of epitope-based diagnostics and vaccines. BMC Bioinformatics. 2007 Dec;8(1):361.

34. Azmi F, Ahmad Fuaad AAH, Skwarczynski M, Toth I. Recent progress in adjuvant discovery for peptide-based subunit vaccines. Human Vaccines & Immunotherapeutics. 2014 Mar;10(3):778–96.

35. Hoover DM, Wu Z, Tucker K, Lu W, Lubkowski J. Antimicrobial Characterization of Human β-Defensin 3 Derivatives. Antimicrob Agents Chemother. 2003 Sep;47(9):2804– 9.

36. Berman H, Henrick K, Nakamura H. Announcing the worldwide Protein Data Bank. Nat Struct Mol Biol. 2003 Dec;10(12):980–980.

37. Berman HM. The Protein Data Bank. Nucleic Acids Research. 2000 Jan 1;28(1):235–42.

38. Schibli DJ, Hunter HN, Aseyev V, Starner TD, Wiencek JM, McCray PB, et al. The Solution Structures of the Human β-Defensins Lead to a Better Understanding of the Potent Bactericidal Activity of HBD3 against Staphylococcus aureus. Journal of Biological Chemistry. 2002 Mar;277(10):8279–89.

39. Walker JM, editor. The Proteomics Protocols Handbook [Internet]. Totowa, NJ: Humana Press; 2005 [cited 2024 Apr 13]. Available from: http://link.springer.com/10.1385/1592598900

40. Magnan CN, Randall A, Baldi P. SOLpro: accurate sequence-based prediction of protein solubility. Bioinformatics. 2009 Sep 1;25(17):2200–7.

41. Cheng J, Randall AZ, Sweredoski MJ, Baldi P. SCRATCH: a protein structure and structural feature prediction server. Nucleic Acids Research. 2005 Jul 1;33(Web Server):W72–6.

42. Hon J, Marusiak M, Martinek T, Kunka A, Zendulka J, Bednar D, et al. SoluProt: prediction of soluble protein expression in Escherichia coli. Xu J, editor. Bioinformatics. 2021 Apr 9;37(1):23–8.

43. Hebditch M, Carballo-Amador MA, Charonis S, Curtis R, Warwicker J. Protein–Sol: a web tool for predicting protein solubility from sequence. Valencia A, editor. Bioinformatics. 2017 Oct 1;33(19):3098–100.

44. Wishart D, Arndt D, Pon A, Sajed T, Guo AC, Djoumbou Y, et al. T3DB: the toxic exposome database. Nucleic Acids Research. 2015 Jan 28;43(D1):D928–34.

45. Cheng J, Saigo H, Baldi P. Large-scale prediction of disulphide bridges using kernel methods, two-dimensional recursive neural networks, and weighted graph matching. Proteins. 2006 Feb 15;62(3):617–29.

46. Buchan DWA, Jones DT. The PSIPRED Protein Analysis Workbench: 20 years on. Nucleic Acids Research. 2019 Jul 2;47(W1):W402–7.

47. Heo L, Park H, Seok C. GalaxyRefine: protein structure refinement driven by side-chain repacking. Nucleic Acids Research. 2013 Jul 1;41(W1):W384–8.

48. Ko J, Park H, Heo L, Seok C. GalaxyWEB server for protein structure prediction and refinement. Nucleic Acids Research. 2012 Jul 1;40(W1):W294–7.

49. Seok C, Baek M, Steinegger M, Park H, Lee GR, Won J. Accurate protein structure prediction: what comes next? BIODESIGN. 2021 Sep 30;9(3):47–50.

50. Laskowski RA, MacArthur MW, Moss DS, Thornton JM. PROCHECK: a program to check the stereochemical quality of protein structures. J Appl Crystallogr. 1993 Apr 1;26(2):283–91.

51. Wiederstein M, Sippl MJ. ProSA-web: interactive web service for the recognition of errors in three-dimensional structures of proteins. Nucleic Acids Research. 2007 May 8;35(Web Server):W407–10.

52. Desta IT, Porter KA, Xia B, Kozakov D, Vajda S. Performance and Its Limits in Rigid Body Protein-Protein Docking. Structure. 2020 Sep;28(9):1071–1081.e3.

53. Vajda S, Yueh C, Beglov D, Bohnuud T, Mottarella SE, Xia B, et al. New additions to the C lus P ro server motivated by CAPRI. Proteins. 2017 Mar;85(3):435–44.

54. Kozakov D, Hall DR, Xia B, Porter KA, Padhorny D, Yueh C, et al. The ClusPro web server for protein–protein docking. Nat Protoc. 2017 Feb;12(2):255–78.

55. Kozakov D, Beglov D, Bohnuud T, Mottarella SE, Xia B, Hall DR, et al. How good is automated protein docking? Proteins. 2013 Dec;81(12):2159–66.

56. Jin MS, Kim SE, Heo JY, Lee ME, Kim HM, Paik SG, et al. Crystal Structure of the TLR1-TLR2 Heterodimer Induced by Binding of a Tri-Acylated Lipopeptide. Cell. 2007 Sep;130(6):1071–82.

57. Park BS, Song DH, Kim HM, Choi BS, Lee H, Lee JO. The structural basis of lipopolysaccharide recognition by the TLR4–MD-2 complex. Nature. 2009 Apr;458(7242):1191–5.

58. Laskowski RA, Jabłońska J, Pravda L, Vařeková RS, Thornton JM. PDBsum: Structural summaries of PDB entries. Protein Science. 2018 Jan;27(1):129–34.

59. Abraham MJ, Murtola T, Schulz R, Páll S, Smith JC, Hess B, et al. GROMACS: High performance molecular simulations through multi-level parallelism from laptops to supercomputers. SoftwareX. 2015 Sep;1–2:19–25.

60. Zoete V, Cuendet MA, Grosdidier A, Michielin O. SwissParam: A fast force field generation tool for small organic molecules. J Comput Chem. 2011 Aug;32(11):2359–68.

61. Ke Q, Gong X, Liao S, Duan C, Li L. Effects of thermostats/barostats on physical properties of liquids by molecular dynamics simulations. Journal of Molecular Liquids. 2022 Nov;365:120116.

62. Rapin N, Lund O, Bernaschi M, Castiglione F. Computational Immunology Meets Bioinformatics: The Use of Prediction Tools for Molecular Binding in the Simulation of the Immune System. Brusic V, editor. PLoS ONE. 2010 Apr 16;5(4):e9862.

63. Stolfi P, Castiglione F, Mastrostefano E, Di Biase I, Di Biase S, Palmieri G, et al. In-silico evaluation of adenoviral COVID-19 vaccination protocols: Assessment of immunological memory up to 6 months after the third dose. Front Immunol. 2022 Oct 24;13:998262.

64. Ragone C, Manolio C, Cavalluzzo B, Mauriello A, Tornesello ML, Buonaguro FM, et al. Identification and validation of viral antigens sharing sequence and structural homology with tumor-associated antigens (TAAs). J Immunother Cancer. 2021 May;9(5):e002694.

65. Madeira F, Pearce M, Tivey ARN, Basutkar P, Lee J, Edbali O, et al. Search and sequence analysis tools services from EMBL-EBI in 2022. Nucleic Acids Research. 2022 Jul 5;50(W1):W276–9.

66. Grote A, Hiller K, Scheer M, Munch R, Nortemann B, Hempel DC, et al. JCat: a novel tool to adapt codon usage of a target gene to its potential expression host. Nucleic Acids Research. 2005 Jul 1;33(Web Server):W526–31.

67. Salim HMW, Cavalcanti ARO. Factors influencing codon usage bias in genomes. J Braz Chem Soc [Internet]. 2008 [cited 2024 Mar 23];19(2). Available from: http://www.scielo.br/scielo.php?script=sci_arttext&pid=S0103-50532008000200008&lng=en&nrm=iso&tlng=en

68. Krebs BB, De Mesquita JF. Amyotrophic Lateral Sclerosis Type 20 - In Silico Analysis and Molecular Dynamics Simulation of hnRNPA1. Xia XG, editor. PLoS ONE. 2016 Jul 14;11(7):e0158939.

69. Knapp B, Frantal S, Cibena M, Schreiner W, Bauer P. Is an Intuitive Convergence Definition of Molecular Dynamics Simulations Solely Based on the Root Mean Square Deviation Possible? Journal of Computational Biology. 2011 Aug;18(8):997–1005.

70. Martínez L. Automatic Identification of Mobile and Rigid Substructures in Molecular Dynamics Simulations and Fractional Structural Fluctuation Analysis. Kleinjung J, editor. PLoS ONE. 2015 Mar 27;10(3):e0119264.

71. Kumar CV, Swetha RG, Anbarasu A, Ramaiah S. Computational Analysis Reveals the Association of Threonine 118 Methionine Mutation in PMP22 Resulting in CMT-1A. Advances in Bioinformatics. 2014 Oct 20;2014:1–10.

72. CDC. Reasons to Get Vaccinated [Internet]. 2023. Available from: https://www.cdc.gov/dengue/vaccine/parents/reasons-to-vaccinate.html

73. Izmirly AM, Alturki SO, Alturki SO, Connors J, Haddad EK. Challenges in Dengue Vaccines Development: Pre-existing Infections and Cross-Reactivity. Front Immunol. 2020 Jun 16;11:1055.

74. Sunita, Sajid A, Singh Y, Shukla P. Computational tools for modern vaccine development. Human Vaccines & Immunotherapeutics. 2020 Mar 3;16(3):723–35.

75. Ferris LK, Mburu YK, Mathers AR, Fluharty ER, Larregina AT, Ferris RL, et al. Human β-Defensin 3 Induces Maturation of Human Langerhans Cell–Like Dendritic Cells: An Antimicrobial Peptide that Functions as an Endogenous Adjuvant. Journal of Investigative Dermatology. 2013 Feb;133(2):460–8.

76. Sormanni P, Aprile FA, Vendruscolo M. The CamSol Method of Rational Design of Protein Mutants with Enhanced Solubility. Journal of Molecular Biology. 2015 Jan;427(2):478–90.

77. Creighton TE. Disulfide bonds as probes of protein folding pathways. In: Methods in Enzymology [Internet]. Elsevier; 1986 [cited 2024 Apr 13]. p. 83–106. Available from: https://linkinghub.elsevier.com/retrieve/pii/007668798631036X

78. Zhang T, Bertelsen E, Alber T. Entropic effects of disulphide bonds on protein stability. Nat Struct Mol Biol. 1994 Jul;1(7):434–8.

79. Woolfson DN. Understanding a protein fold: The physics, chemistry, and biology of α-helical coiled coils. Journal of Biological Chemistry. 2023 Apr;299(4):104579.

